# Nitrate-dependent salt tolerance mediated by OsNLP4-OsMADS27 module

**DOI:** 10.1101/2022.07.08.499223

**Authors:** Alamin Alfatih, Jing Zhang, Ying Song, Sami Ullah Jan, Zi-Sheng Zhang, Jing-Qiu Xia, Zheng-Yi Zhang, Tahmina Nazish, Jie Wu, Ping-Xia Zhao, Cheng-Bin Xiang

## Abstract

Salt stress is a major constraint of plant growth and yield. Nitrogen (N) fertilizers are known to alleviate salt stress. However, the underlying molecular mechanisms remain unclear. Here we show that OsNLP4-OsMADS27 module controls nitrate-dependent salt tolerance in rice. The expression of *OsMADS27* is specifically induced by nitrate. The *OsMADS27* knockout mutants are more sensitive to salt stress than the wild type, whereas the *OsMADS27* overexpression lines are more tolerant. Transcriptomic analyses revealed that OsMADS27 controls the expression of a number of known stress-responsive genes as well as those involved in ion homeostasis and antioxidation. We demonstrated that OsMADS27 directly binds to the promoter of *OsHKT1.1* and *OsSPL7* to regulate their expression. Notably, *OsMADS27*-mediated salt tolerance is nitrate-dependent and positively correlated with nitrate concentration. We further showed that OsNLP4, a nitrate-responsive key regulator in N metabolism and N use efficiency, positively regulates the expression of *OsMADS27* by directly binding to the nitrate-responsive *cis*-element in its promoter, thereby transmitting the nitrate signal to *OsMADS27* and conferring its nitrate dependence. Our results reveal the role of nitrate-responsive OsNLP4-OsMADS27 module and its downstream target genes in salt tolerance, filling the gap in the molecular mechanism of nitrate-dependent salt tolerance of rice. Moreover, *OsMADS27* overexpression increased grain yield under salt stress in presence of sufficient nitrate, indicating that *OsMADS27* is a promising candidate for the improvement of salt tolerance in rice.

## Introduction

Salinity lies among critical crises in agriculture around the globe and the majority of the food crops are salinity-sensitive (Qadir et al., 2014). Elevated soil salinity not only causes ion toxicity and osmotic stress, but also results in severe nutrient deficiency in plants (Munns and Tester, 2008). To cope with the salinity-triggered damages, plants have evolved various strategies on the bases of their habitat and severity of stress (Adem et al., 2014; Ashraf et al., 2008; Bose et al., 2014; Chakraborty et al., 2016). Among numerous strategies, appropriate acquisition of the mineral nutrients is undoubtedly an effective way to improve salinity tolerance, growth and yield under salt stress (Gao et al., 2016; Guo et al., 2017; Kaya et al., 2007). Therefore, it is important to understand the mechanisms by which nutrients alleviate salt stress in plants for breeding robust salt-tolerant crop varieties.

Potassium, a vital nutrient for plant growth and development, is well known for its role in balancing sodium concentration in plants (Clarkson and Hanson, 1980; Raddatz et al., 2020; Wu et al., 2018; Zorb et al., 2014). Under salt stress, the accumulation of sodium ions (Na^+^) in the cytoplasm leads to membrane depolarization and promotes potassium ions (K^+^) leakage out of the cell. Therefore, it is crucial for plants to maintain an appropriate K^+^/Na^+^ ratio in the cytoplasm to survive in saline soil, which depends on the operation of Na^+^/ K^+^ transporters (Wu et al., 2018). Rice potassium transporter OsHAK1 promotes K^+^ uptake and K^+^/Na^+^ ratio in both low and high potassium conditions, which is essential for maintaining potassium-mediated growth and salt tolerance (Chen et al., 2015). Rice shaker K^+^ channel OsAKT2 mediates K^+^ recirculation from shoots to roots to maintain Na^+^/K^+^ homeostasis and improve salt tolerance (Tian et al., 2021). Moreover, the members of high-affinity K^+^ transporters like HKTs also grant salinity tolerance to rice (Hamamoto et al., 2015; Rosas-Santiago et al., 2015; Suzuki et al., 2016a; Wang et al., 2015). Calcium (Ca^2+^) can regulate the perception, uptake, and transport of various ions through the SOS (salt overly sensitive) pathway (Lin et al., 2009; Qiu et al., 2002; Yang and Guo, 2018a; Yang and Guo, 2018b; Zhu et al., 1998), thereby coordinating Na^+^/K^+^ homeostasis in plants (Asano et al., 2012; Campo et al., 2014; Manishankar et al., 2018). The Na^+^/H^+^ antiporter SOS1 in cell membrane is associated with Na^+^ extrusion via roots under saline environment and confers salinity tolerance to rice (Martínez-Atienza et al., 2007). *SOS2* and *SOS3*, encoding protein kinase and Ca^2+^-binding protein respectively, are required for salinity tolerance in rice because they perceive the change of Ca^2+^ in cytosol under salinity and activate several downstream genes to start signaling cascade (Kumar et al., 2013).

Apart from potassium, few mineral nutrients have been studied for their roles in salt tolerance. Sulfur nutrient has been found to improve plant photosynthesis and growth under salt stress by increasing glutathione production and abscisic acid (ABA) accumulation (Cao et al., 2014; Chen et al., 2019; Fatma et al., 2014; Fatma et al., 2021). Nitrogen (N), an essential macronutrient for plants, has been shown to improve salt tolerance (Mansour, 2000) via its participation in stimulation of the antioxidation (Rais et al., 2013), osmotic adjustment (Nasab et al., 2014), maintenance of ion balance (Khan et al., 2016b), mitigation of ionic toxicity (Iqbal et al., 2015), and the activation of numerous enzymes (Aragao et al., 2012). However, the underlying molecular mechanisms of N-improved salt tolerance in plants remain unclear to date.

Transcription factors (TFs) play essential roles in transcriptional control of the stress-associated genes and hence are of utmost importance for breeding stress-tolerant crops (Ahammed et al., 2020; Zhang et al., 2017). The MADS family TFs control important growth and developmental processes such as seed germination and flowering time (Chen et al., 2016; Moyle et al., 2005; Wu et al., 2017; Yin et al., 2019; Yu et al., 2017). MADS-box TFs are also involved in the response to various abiotic stress. For example, *OsMADS26* is a negative regulator of drought stress tolerance in rice (Khong et al., 2015). *OsMADS57* in concert with *OsTB1* mediates the transcription of *OsWRKY94* to confer cold tolerance in rice (Chen et al., 2018b). Moreover, *OsMADS25*, *OsMADS27* and *OsMADS57* are involved in the response to nutrient deficiency in rice (Chen et al., 2018a; Huang et al., 2019; Yu et al., 2015). The overexpression of *OsMADS25* improved the salinity tolerance of rice (Wu et al., 2020).

We previously reported *Arabidopsis* MADS-box TF *AtAGL16* as a negative regulator of salt and drought tolerance (Zhao et al., 2020; Zhao et al., 2021). To extend our work to rice, we identified *OsMADS27* as the most homologous gene of *AtAGL16*. *OsMADS27* is induced by nitrate (NO_3_^-^) and ABA, and acts as a target gene of miR444 to control root development in a NO_3_^-^-dependent manner (Chen et al., 2018a; Pachamuthu et al., 2022; Puig et al., 2013; Yu et al., 2014). When overexpressed, *OsMADS27* confers enhanced salt tolerance in transgenic seedlings (Chen et al., 2018a). However, the molecular mechanism underlying *OsMADS27*-mediated salt tolerance remains unclear. Likewise, the relation of *OsMADS27*-mediated salt tolerance to N nutrient has not been investigated in rice. In this study, we discovered the NO_3_^-^ dependence of *OsMADS27*-mediated salt tolerance and unveiled the underlying molecular mechanism. Our results demonstrate that *OsMADS27*-mediated salt tolerance is NO_3_^-^-dependent. The OsNLP4-OsMADS27 module plays a key role in the NO_3_^-^-dependent salt tolerance, where OsNLP4 senses NO_3_^-^ signaling, translocates to the nucleus (Konishi and Yanagisawa, 2010; Wu et al., 2021), and transcriptionally upregulates *OsMADS27*. Consequently, OsMADS27 directly regulates the expression of stress-responsive genes in rice. Therefore, our findings revealed a novel mechanism of NO_3_^-^-dependent salt tolerance, which can be exploited for the improvement of salinity tolerance in crops.

## Results

### Expression of *OsMADS27* is specifically induced by nitrate and NaCl-induced expression of *OsMADS27* is nitrate-dependent

To gain a detailed expression pattern of *OsMADS27*, we examined its spatiotemporal expression by quantitative real time PCR (qRT-PCR) at three developmental stages of rice plants: seedling, vegetative, and pre-mature stage. Our results demonstrated that *OsMADS27* was expressed in all the tissues examined but with much higher levels in roots, leaves, and sheath (Fig. S1A). In addition, tissue expression pattern of *OsMADS27* was revealed in the *OsMADS27pro*::*GUS* transgenic plants (Fig. S1B), which was reconcilable with our qRT-PCR results and previous reports (Chen et al., 2018a; Pachamuthu et al., 2022; Puig et al., 2013; Yu et al., 2014). Notably, strong GUS signal was detected in the stelle of the root (Fig. S1B).

To check the response of *OsMADS27* to nutrients and salt stress, we grew wild type (WT) seedlings under normal conditions and then transferred 7-day-old seedlings to hydroponic medium without N for 48 hours, then transferred the seedlings to hydroponic medium supplemented with 2 mM KNO_3_, 2 mM NH_4_Cl, 2 mM KCl, or 150 mM NaCl, respectively. Surprisingly, NaCl did not induce the expression of *OsMADS27* under our conditions, neither did NH_4_Cl or KCl. Only KNO_3_ induced the expression of *OsMADS27* that plateaued at 12 hours with about 5 folds increase (Fig. 1A). In addition, we showed that when the seedlings were transferred into N-free medium, the KNO_3_-induced expression of *OsMADS27* was gradually decreased (Fig. 1B). These results clearly show that the expression of *OsMADS27* is specifically responsive to KNO_3_.

**Figure 1.**
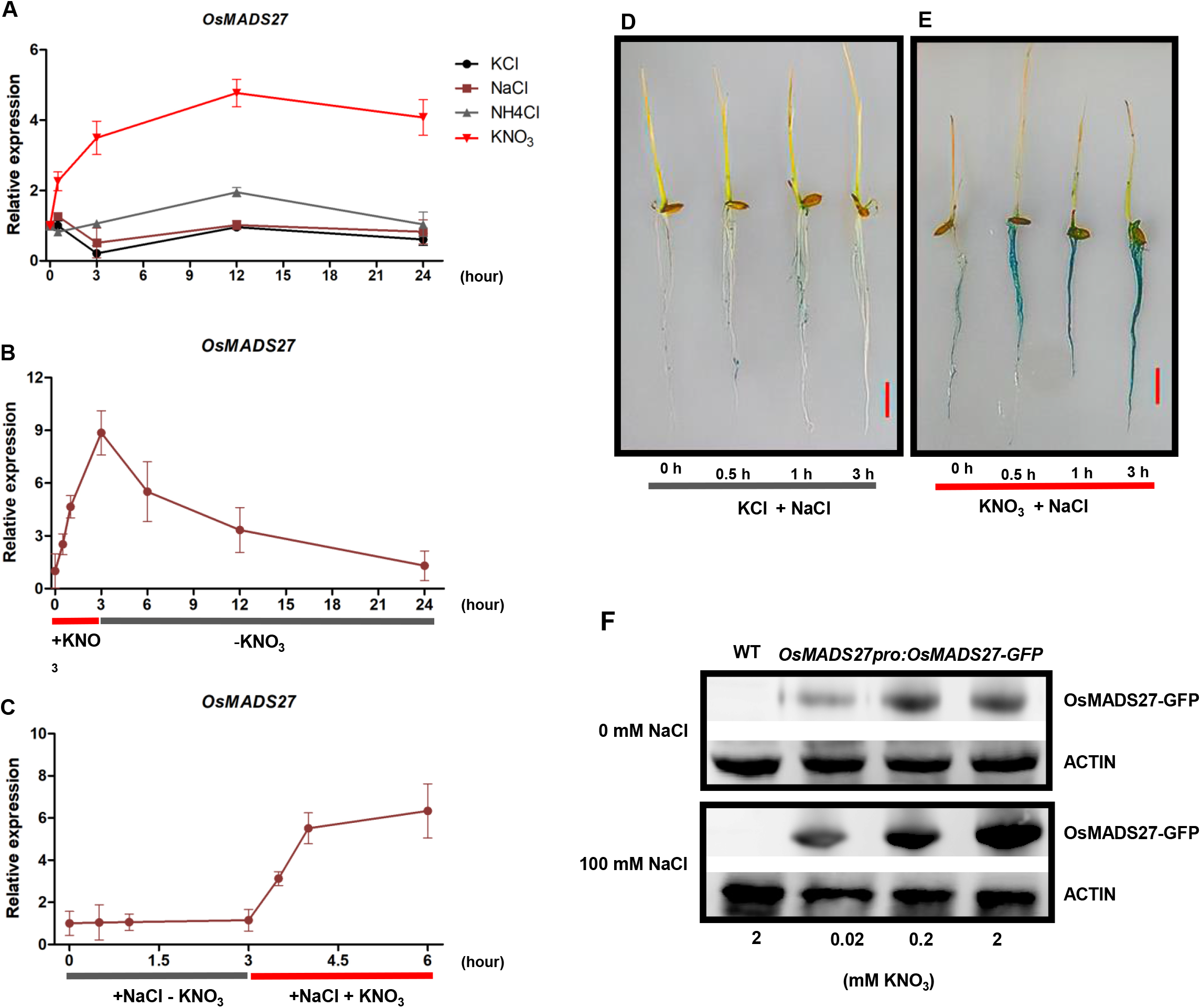
*OsMADS27* is specifically responsive to nitrate. A. Time-course analyses of *OsMADS27* expression in response to N and salt stress. 7-day-old wild type plants grown on hydroponic medium with 1.5 mM KNO_3_ were transferred to hydroponic medium without N for 2 days, and then transferred to hydroponic medium with 2 mM KNO_3_, 2 mM NH_4_Cl, 140 mM NaCl or 2 mM KCl for 0, 0.5, 3, 12, 24 hours. RNA was extracted from the whole seedlings for qRT-PCR analyses as described in the Materials and method. Values are mean ± SD (n = 3). B. Time-course analyses of *OsMADS27* expression in response to KNO_3_ depletion. 7-day-old wild type plants grown under on hydroponic medium with 1.5 mM KNO_3_ were treated with 2 mM KNO_3_ for 0, 0.5, 1, 3 hours, then transferred to hydroponic medium without KNO_3_ for 3, 6, 12, 24 hours. RNA was extracted from the whole seedlings for qRT-PCR analyses as described in the Materials and method. Values are mean ± SD (n = 3). C. KNO_3_-dependent induction of *OsMADS27* expression by NaCl. Wild type seedlings hydroponically grown on N-free medium for 7 days were treated with 140 mM NaCl for 0, 0.5, 1, 3 hours, and then transferred to hydroponic medium with 140 mM NaCl + 2 mM KNO_3_ for 0.5, 1, 3 hours. RNA was extracted from the whole seedlings for qRT-PCR analyses as described in the Materials and method. Values are mean ± SD (n = 3). D-E. The response of *OsMADS27pro*:*GUS* to NaCl. 7-day-old *OsMADS27pro*:*GUS* lines grown on N-free medium with 2 mM KCl (D) or 2 mM KNO_3_ (E) were treated with 140 mM NaCl for 0.5, 1, 3 hours, respectively. Seedlings were incubated in GUS buffer for 3 hours before photographs were taken. Bar = 1 cm F. OsMADS27 protein level in *OsMADS27pro:OsMADS27-GFP* plants. 2-week-old *OsMADS27pro :OsMADS27-GFP* seedlings grown hydroponically on medium contains different N concentrations (0.02 mM, 0.2 mM and 2 mM KNO_3_) without (control) or with 100 mM NaCl were used for the analysis of OsMADS27 protein level by western blot with anti-GFP antibodies. ZH11 (WT) grown on medium with 2 mM KNO_3_ served as a control.

Meanwhile, we showed that under normal growth conditions, NaCl and ABA induced the expression of *OsMADS27* (Fig. S1C-E), which was inconsistent with the results of NaCl treatment in Fig. 1A. The only difference of these experiments lies in whether nitrate is present in the NaCl treatment, which likely counts for this observed difference of *OsMADS27* expression. To confirm this, we treated seedling (N starved) under 150 mM NaCl with 0 mM KNO_3_ for 3 hours, then added 2 mM KNO_3_ for another 3 hour. The qRT-PCR results clearly show that in the absence of KNO_3_, NaCl was unable to induce the expression of *OsMADS27*. Only in the presence of KNO_3_, NaCl stimulated the expression of *OsMADS27* (Fig. 1C). This was further confirmed with *OsMADS27pro:GUS* transgenic rice in which GUS signal exhibited a similar response. No change in GUS activity was observed under the treatment of KCl plus NaCl, while a strong induction of GUS was seen in roots treated with KNO_3_ plus NaCl (Fig. 1D and E).

We also quantified the protein level of OsMADS27 in the *OsMADS27pro:OsMADS27-GFP* plants by western blot using anti-GFP antibodies under low, normal, and high concentration of KNO_3_ (0.02 mM, 0.2 mM, and 2mM) with or without 100 mM of NaCl for 10 days. The results in Fig. 1F show that the OsMADS27 protein level is positively correlated with KNO_3_ concentration and enhanced by NaCl treatment (Fig. 1F).

Taken together, our results clearly show that the expression of *OsMADS27* is specifically induced by NO_3_^-^ and NaCl-induced expression of *OsMADS27* is NO_3_^-^-dependent.

### Nuclear localization of OsMADS27 is responsive to nitrate

To reveal the subcellular localization of OsMADS27 protein and its response to nutrients, we generated *OsMADS27pro:OsMADS27-GFP* transgenic lines. The transgenic plants were grown on N-free MS medium supplied with 2 mM KNO_3_ (Fig. 2A) or 2 mM KCl (Fig. 2C) for 10 days. Then seedlings receiving KNO_3_ were treated with 150 mM NaCl (Fig. 2B), and the seedlings receiving KCl were treated with 2 mM KNO_3_ (Fig. 2D), 2 mM NH_4_Cl (Fig. 2E), 150 mM NaCl (Fig. 2F), and 150 mM NaCl plus 2 mM KNO_3_ (Fig. 2G) respectively for 60 min before confocal laser-scanning microscopic observation. GFP signals were detected in the nucleus whenever KNO_3_ was included in the medium regardless of the presence of other supplements (Fig. 2A, B, D and G). No GFP signals were detected in the presence of KCl (Fig. 2C), KCl plus NH_4_Cl (Fig. 2E), or KCl plus NaCl (Fig. 2F). These results indicate that the nuclear accumulation of OsMADS27 is specifically responsive to nitrate, in accordance with that of OsNLP4 (Wu et al., 2021).

**Figure 2.**
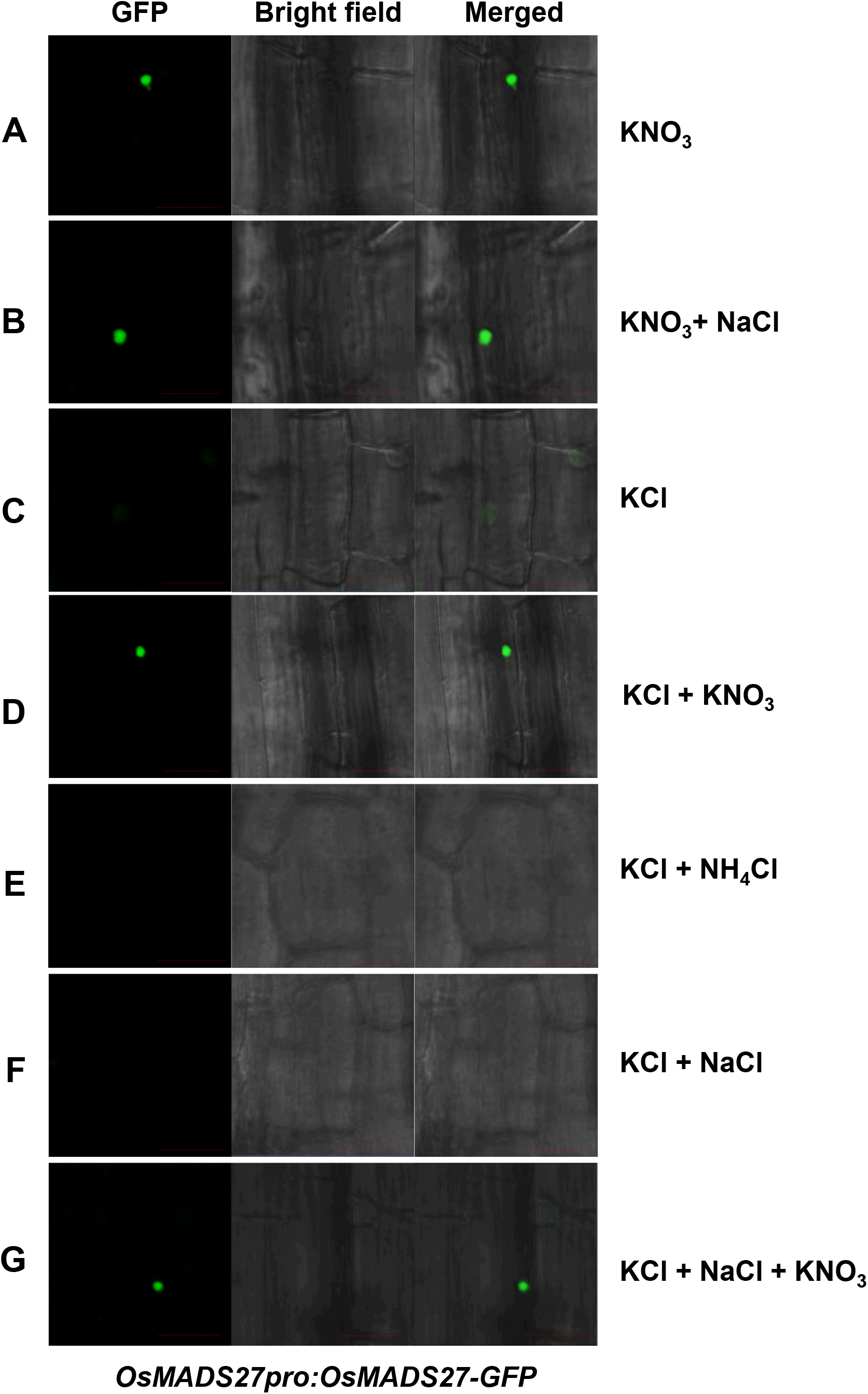
Nitrate-responsive nuclear localization of OsMADS27. *OsMADS27pro:OsMADS27-GFP* plants were grown on N-free MS medium supplied with 2 mM KNO_3_ (A) or 2 mM KCl (C) for 10 days. The seedlings in A were treated with 150 mM NaCl (B) for 60 min before green fluorescence observation. The seedlings in C were treated with 2 mM KNO_3_ (D), 2 mM NH_4_Cl (E), 150 mM NaCl (F), and 150 mM NaCl + 2 mM KNO_3_ (G) respectively for 60 min before GFP observation. The green fluorescence was observed on the Zeiss 880 microscope. Scale bars = 20 μm.

### *OsMADS27*-mediated salt tolerance in rice is nitrate-dependent

To explore the capability of *OsMADS27* in conferring salt tolerance to rice, we generated two independent loss-of-function mutant lines of *OsMADS27* (KO1 and KO2) by using CRISPR/CAS9-based editing. Protein sequence alignment depicted that mutations in both mutants resulted in premature stop codon, hence interrupting the open reading frame (ORF) of *OsMADS27* (Fig. S2A-D). Additionally, we generated two independent overexpression (OE) lines of *OsMADS27* (OE7 and OE8) driven by *OsACTIN1* promoter (Fig. S2E-F).

To evaluate the role of *OsMADS27* in salt stress tolerance of rice, we germinated the seeds of OE7, OE8, KO1, KO2, and WT in soil in the presence of 0 mM or 150 mM NaCl. Under 0 mM NaCl conditions, there was no difference in germination rate among all the genotypes (Fig. S3A). However, under 150 mM salt stress, OE lines displayed a germination rate of 80% at day 6 compared with WT and KO mutants which exhibited a germination rate of 55% and 30% respectively (Fig. S3B). Moreover, we conducted salt tolerance assay on soil-grown seedlings (Fig. S3C). Upon treatment of 20-day-old soil-grown seedlings with 150 mM NaCl for 15 days, 80% of the OE plants survived compared with WT and KO mutants with a survival ratio of 43% and 12% respectively, whereas under the 0 mM NaCl control treatment all genotypes displayed 100% survival (Fig. S3D). Together these results clearly demonstrate that *OsMADS27* is a positive regulator of salt tolerance in rice.

The NO_3_^-^ dependence of NaCl-induced expression of *OsMADS27* prompted us to ask whether OsMADS27-mediated salt tolerance is nitrate-dependent. Thus we further explored the salt tolerance of different *OsMADS27* genotypes under different NO_3_^-^ concentrations. We grew seedlings in modified hydroponic culture with different NO_3_^-^ concentrations for 10 days, then supplemented with or without 140 mM NaCl in the hydroponic culture and allowed seedlings to grow for another week. Under 0 mM NaCl conditions, the seedling survival rate was 100% for all the genotypes under all three concentrations of NO_3_^-^ (Fig. 3A and C). Under 140 mM NaCl conditions, the seedling survival rate of all the three genotypes was similarly less than 20% under low NO_3_^-^ conditions (0.02 mM, LN). However, under normal NO_3_^-^ conditions (0.2 mM, NN), the increased NO_3_^-^ alleviated the salt stress as reflected by the seedling survival rate of KO mutants (20%), WT (42%) and OE lines (55%) compared with those under LN conditions (Fig. 3B and D). Under high NO_3_^-^ conditions (2 mM, HN), salt stress was further alleviated as evidenced by the increased seedling survival rate in KO mutants (30%), WT (70%), and OE lines (80%). These results demonstrate that the salt tolerance mediated by *OsMADS27* is NO_3_^-^-dependent.

**Figure 3.**
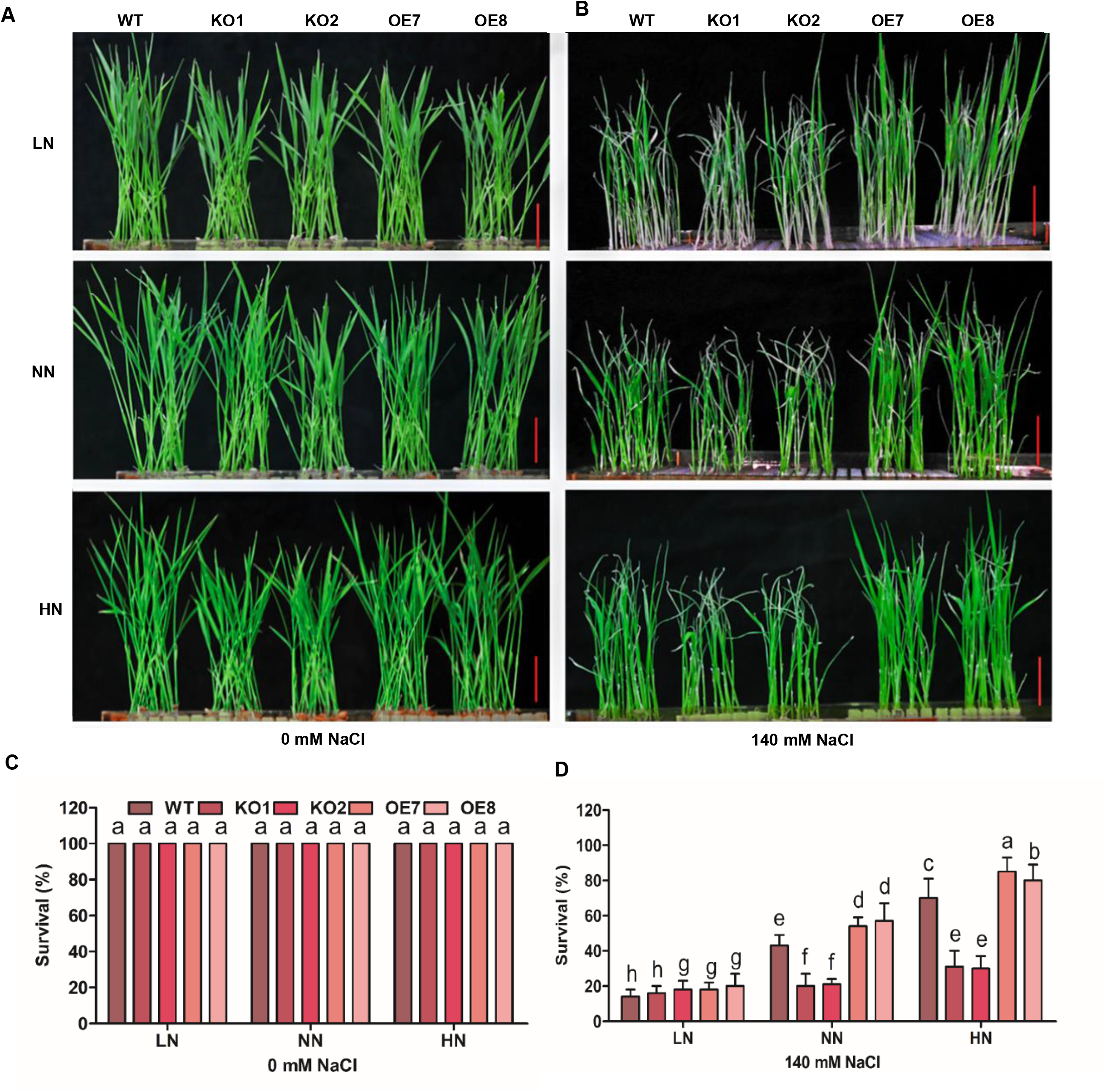
Nitrate-dependent salt tolerance of seedlings. A-D. Hydroponic salt tolerance assay. Seeds of WT (ZH11), KO1, KO2 mutants, OE7 and OE8 lines were germinated at 37 °C for 4 days and transferred to modified hydroponic medium containing different N concentration (0.02 mM, 0.2 mM, 2 mM KNO_3_) for 7 days followed by application of 0 mM, 140 mM NaCl for 7 days before photographs were taken (A-B), and survival rate was calculated (C-D). Values are mean ± SD (n=3 replicates, 32 seedlings per replicate).

To confirm the above hydroponic culture results, we grew the plants of the three genotypes in potted vermiculite and fed with nutrient solutions containing different concentrations of NO_3_^-^ (1.5 mM LN, 2.5 mM NN, 5 mM HN) with or without 65 mM NaCl as described in Methods (Fig. S4A). The data of yield-related agronomic traits were collected for statistical analyses. Fig. S4B shows the grain yield per plant of the three genotypes under three N levels without salt stress. The OE line exhibited significantly higher yield than WT at all three N levels, while the KO showed lower yield than WT. The OE line exhibited grain yield increase by 29%, 38%, and 25% relative to the WT under LN, NN, and HN conditions respectively, while the KO dispayed yield decrease by 20%, 22%, and 25%. The yield was positively correlated with N level, tiller number per plant (Fig. S4C), and panicle number (Fig. S4D). Both tiller and panicle number dispayed similar pattern of genotype and N level effects as grain yield did. Under salt stress, The OE line exhibited grain yield increase by 66%, 40%, and 28% relative to the WT under LN, NN, and HN conditions respectively, while the KO dispayed yield decrease by 33%, 40%, and 28% under the same conditions (Fig. S4E). Tiller and panicle number displayed a similar trend as grain yield did (Fig. S4F and G). These results suggest that *OsMADS27* is a positive regulator of grain yield and further support that *OsMADS27* positively regulates salt tolerance in a NO_3_^-^-dependent manner in rice.

We also conducted field trials to examine the yield of three *OsMADS27* genotypes in the field of varying N supply and found that agronomic traits including nitrogen use efficiency (NUE), actual yield per plot, grain yield per plant, panicles number per plant, number of seeds per plant, and primary branch number per panicle were significantly improved in OE plants under normal and high N availability, whereas reduced in KO plants compared with the wild type (Fig. S5). The field trial data further support that *OsMADS27* is a positive regulator of grain yield, which is positively correlated with NO_3_^-^ availability.

### RNA sequencing reveals *OsMADS27-*regulated genes involved in stress tolerance

To determine the global network of genes regulated by *OsMADS27*, we carried out transcriptomic analyses of WT, KO, and OE plants subjected to 0 mM or 100 mM NaCl for 3 consecutive days to identify the DEGs (differentially expressed genes). The number of DEGs was significantly different among WT, KO, and OE under saline and normal conditions, revealing that *OsMADS27* widely regulates the transcriptome in response to salt stress (Fig. 4A and B).

**Figure 4.**
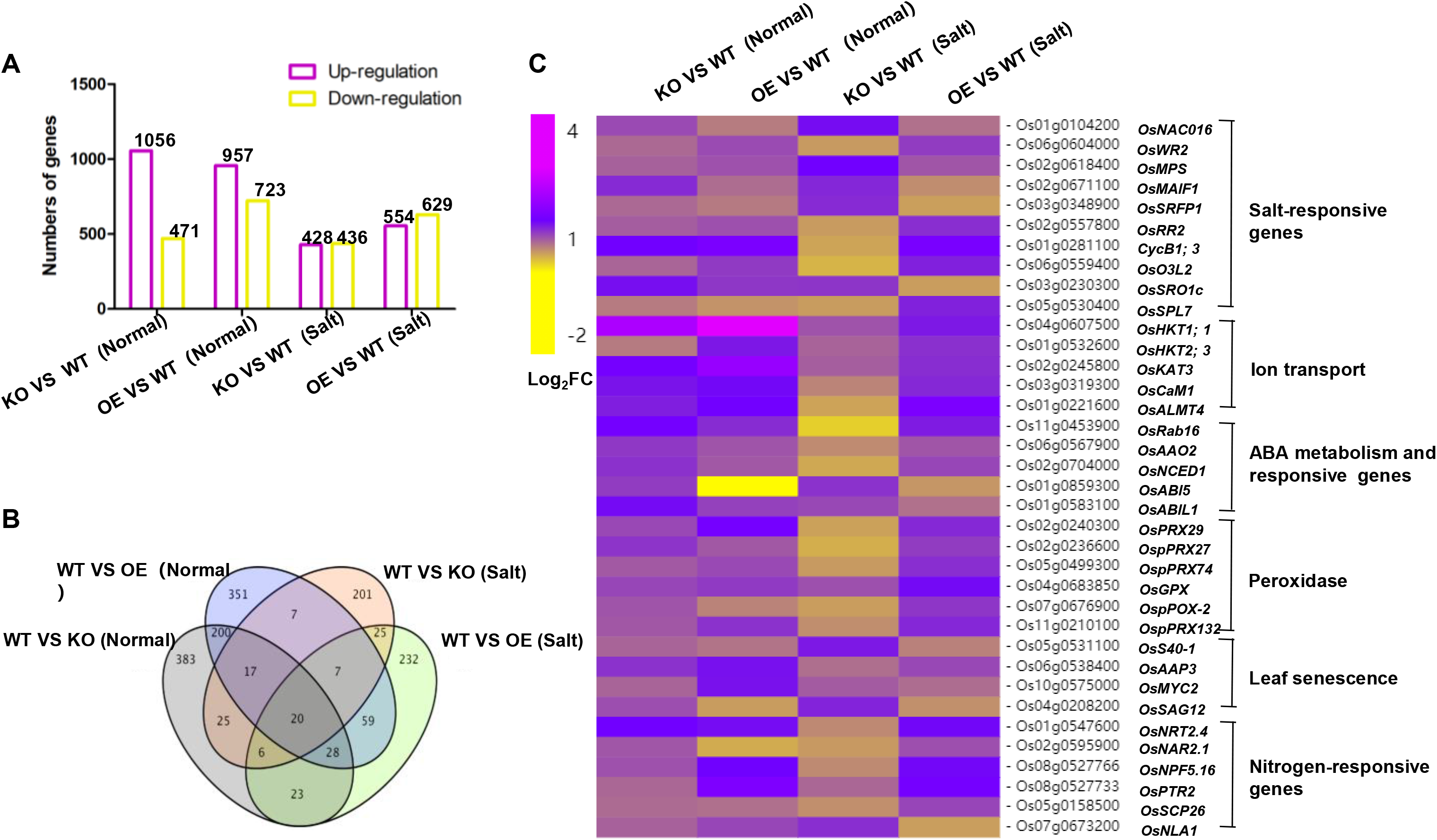
Transcriptomic analysis of differentially expressed genes (DEGs) affected by OsMADS27. A. The number of differentially expressed genes (DEGs). The statistics data of differentially expressed genes in (KO vs WT)-control, (OE vs WT)-control, (KO vs WT)-salt, and (OE vs WT)-salt groups. B. Venn diagram of differentially expressed genes (DEGs) among (KO vs WT)-control, (OE vs WT)-control, (KO vs WT)-salt, and (OE vs WT)-salt groups. The numbers represent the total numbers of differentially expressed genes in different comparison groups. C. Hierarchical clustering analysis of N and salt stress-related genes affected by OsMADS27 in DEGs. The heatmap represents fold changes in the abundance of gene transcripts in different comparison groups.

The in-depth information about DEGs was obtained by KEGG (Kyoto Encyclopedia of Genes and Genomes) pathway and GO (Gene Ontology) analyses to detect significantly expressed DEGs in KO vs WT and OE vs WT under control and salt conditions (Fig. 4C and S6). Remarkably, the genes involved in salt response were highly enriched in DEGs, indicating that *OsMADS27* may coordinately regulate the key genes in salt tolerance (Fig. 4C). The heatmap demonstrates that the transcript level of ethylene response factor (*OsWR2*), salinity-responsive MYB transcription factor (*OsMPS*), A-type response regulator (*OsRR2*), rice cyclin gene (*OsCycB1;3*), oxidative stress 3 (*OsO3L2*), and a heat shock transcription factor (*OsSPL7*) was higher in the OE plants under salt stress. In addition to the salt-responsive genes, key genes involved in ion transport, such as K^+^ transporters (*OsHKT1.1*, *OsHKT2.3*), K^+^ channel (*OsKAT3*), salt-inducible calmodulin gene (*OsCAM1*), and aluminum-activated transporter of malate (*OsALMT4*) were significantly down-regulated in KO mutant while up-regulated in the OE line under salt stress. *OsMADS27* also positively regulates the expression of prominent ABA-responsive genes such as *OsNCED1*, *OsRAB16,* and *OsGLP1*, which were expressed at higher levels in OE plants. Moreover, the genes of peroxidases in antioxidation including *OsPRX29*, *OsPRX27*, *OsPRX74*, *OsGPX*, *OsPRX132* were significantly upregulated in OE vs WT (salt group). Furthermore, *OsMADS27* positively regulates the expression of N-responsive genes as the expression level of *OsNRT2.4*, *OsNAR2.1*, *OsNPF5.16*, *OsNPF2.2*/*OsPTR2* and *OsNLA1* was predominantly enhanced in WT vs OE group after salt treatment (Fig. 4C). In addition, GO enrichment analyses show that *OsMADS27* also affected the expression of some genes involved in oxidation-reduction process, regulation of transcription, defense response and protein phosphorylation under normal conditions (Fig. S6A and B), whereas hydrogen peroxide catabolic process, flavonoid biosynthesis, abscisic acid catabolism, defense response and tyrosine catabolism related genes were also regulated by *OsMADS27* under salt stress conditions (Fig. S6C and D).

The expression pattern of the genes involved in salt response and ion transport was verified by RT-qPCR, which was largely in agreement with the RNA-seq data (Fig. S7). Taken together, our RNA-seq data suggest that *OsMADS27* confers salt tolerance in rice by regulating salt-responsive genes, maintaining ion balance, and enhancing ROS scavenging.

### OsMADS27 transcriptionally activates *OsHKT1.1* and *OsSPL7*

To demonstrate the capability of OsMADS27 to regulate its target genes, we generated the transgenic rice plants expressing *OsMADS27pro:OsMADS27-GFP* for ChIP (chromatin immunoprecipitation) assay. The *cis*1 region of *OsHKT1.1* promoter and *cis*2 and *cis*3 regions of *OsSPL7* promoter were found to be enriched in the transgenic rice plants as demonstrated by qRT-PCR (Fig. 5A and B). Furthermore, we performed transactivation assays using 35S-*OsMADS27* as the effector and *OsHKT1.1* and *OsSPL7* promoter-driven LUC (luciferase) as reporters. When reporter and effector were co-transfected into the tobacco leaves, we observed that *OsMADS27* activated the expression of LUC genes linked to the promoters of *OsHKT1.1* and *OsSPL7* (Fig. 5C and D). Taken together, these results demonstrate that OsMADS27 binds the *cis* elements in the promoter of *OsHKT1.1* and *OsSPL7* and activates their expression.

**Figure 5.**
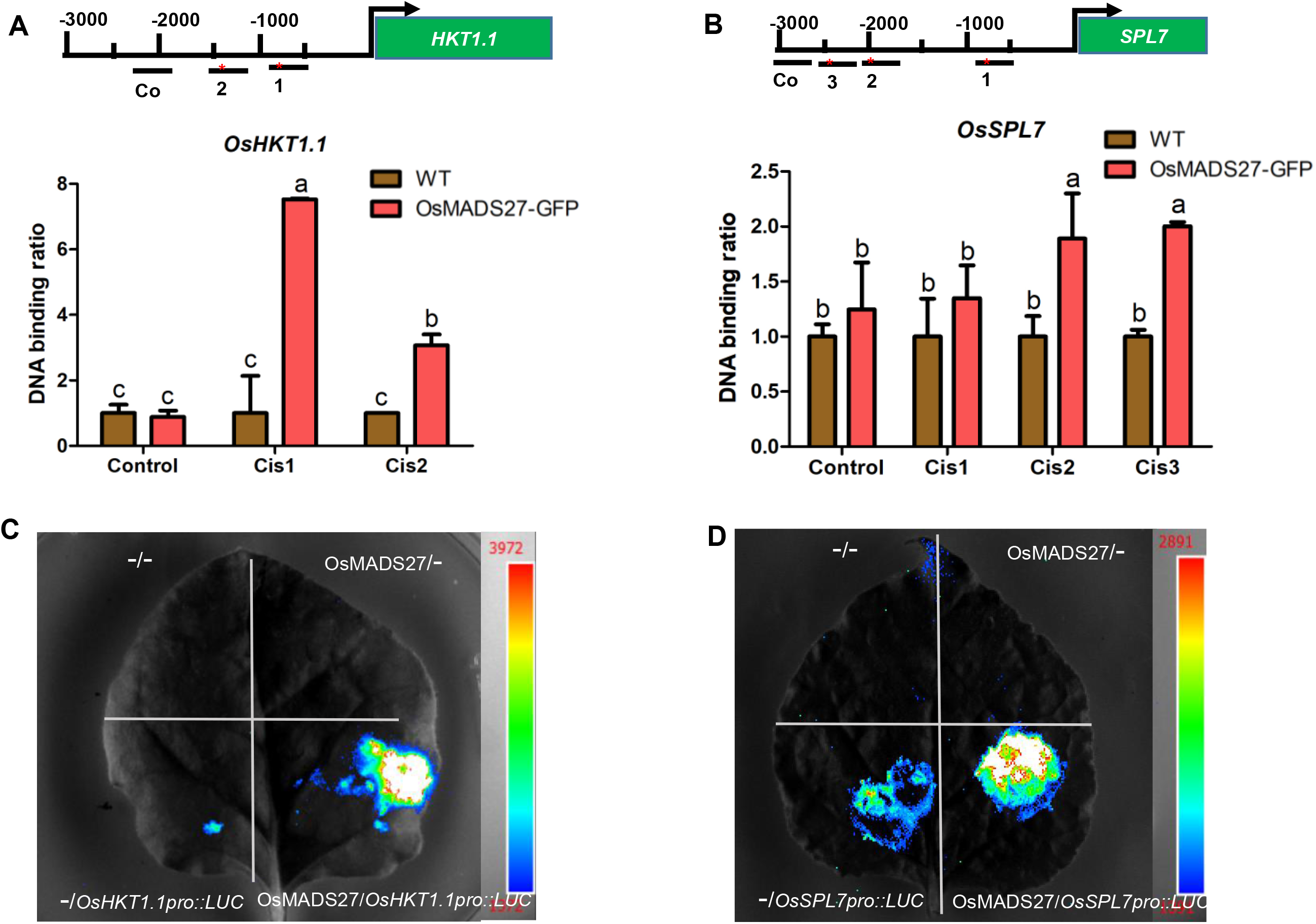
OsMADS27 activates *OsHKT1.1* and *OsSPL7* by binding to the CArG motif in their promoter. A-B. ChIP-qPCR assay. The enrichment of the fragments containing CArG motifs (marked with asterisks) in promoters of *OsHKT1.1* and *OsSPL7* was checked in *OsMADS27pro:OsMADS27-GFP* and wild type plants. About 200 bp fragment *cis*1 and *cis*2 of *OsHKT1.1* promoter (A), *cis*2 and *cis*3 of *OsSPL7* promoter (B) were enriched in *OsMADS27pro:OsMADS27-GFP* plants by anti-GFP antibodies as shown in qRT-PCR analyses. Values are mean ± SD (n=3 replicates). Different letters denote significant differences (P < 0.05) from Duncan’s multiple range tests. C-D. Luciferase activity assay. pRI101-*OsMADS27* acts as effector. pGreenII0800-*OsHKT1.1pro::LUC* / *OsSPL7pro::LUC* function as reporters. “-/-” represents pRI101 and pGreenII 0800 empty plasmids. “-/-”, “OsMADS27/-”, “-/*OsHKT1.1pro::LUC*”, “- /*OsSPL7pro::LUC*” as negative controls; “OsMADS27/*OsHKT1.1pro::LUC*” (E), “OsMADS27/ *OsSPL7pro::LUC*” (F) as experimental groups. Different constructs were separately coinfiltrated into 4-week-old tobacco leaves, then the luciferase activity was detected by the luciferase assay system.

### Nitrate-responsive OsNLP4 upregulates *OsMADS27* and confers its nitrate dependence

To explore the mechanism by which NO_3_^-^ specifically induces the expression of *OsMADS27*, we performed *cis* elements search in the promoter of *OsMADS27* and found that the promoter of *OsMADS27* harbors multiple nitrate-responsive *cis*-elements (NREs), the binding site for nin-like protein (NLP) transcription factors (Konishi and Yanagisawa, 2010; Wu et al., 2021). Further exploration revealed that the expression level of *OsMADS27* was significantly up-regulated in *OsNLP4* overexpression plants and down-regulated in the knockout mutants (Fig. 6A). Subsequent ChIP assay showed that the *cis*1 portion of the *OsMADS27* promoter harboring NRE was significantly enriched, confirming that OsNLP4 binds to the *OsMADS27* promoter *in vivo* (Fig. 6B). The binding was further confirmed by electrophoretic mobility shift assay (EMSA) (Fig. 6C). Furthermore, we conducted a dual-luciferase reporter assay to further verify the transcription activation of *OsMADS27* by OsNLP4 in tobacco leaves. Strong fluorescent signals were shown when the effector construct *35S-OsNLP4* was co-transfected with the reporter construct *OsMADS27pro::LUC* (Fig. 6D), indicating that OsNLP4 transcriptionally activates the expression of *OsMADS27*.

**Figure 6.**
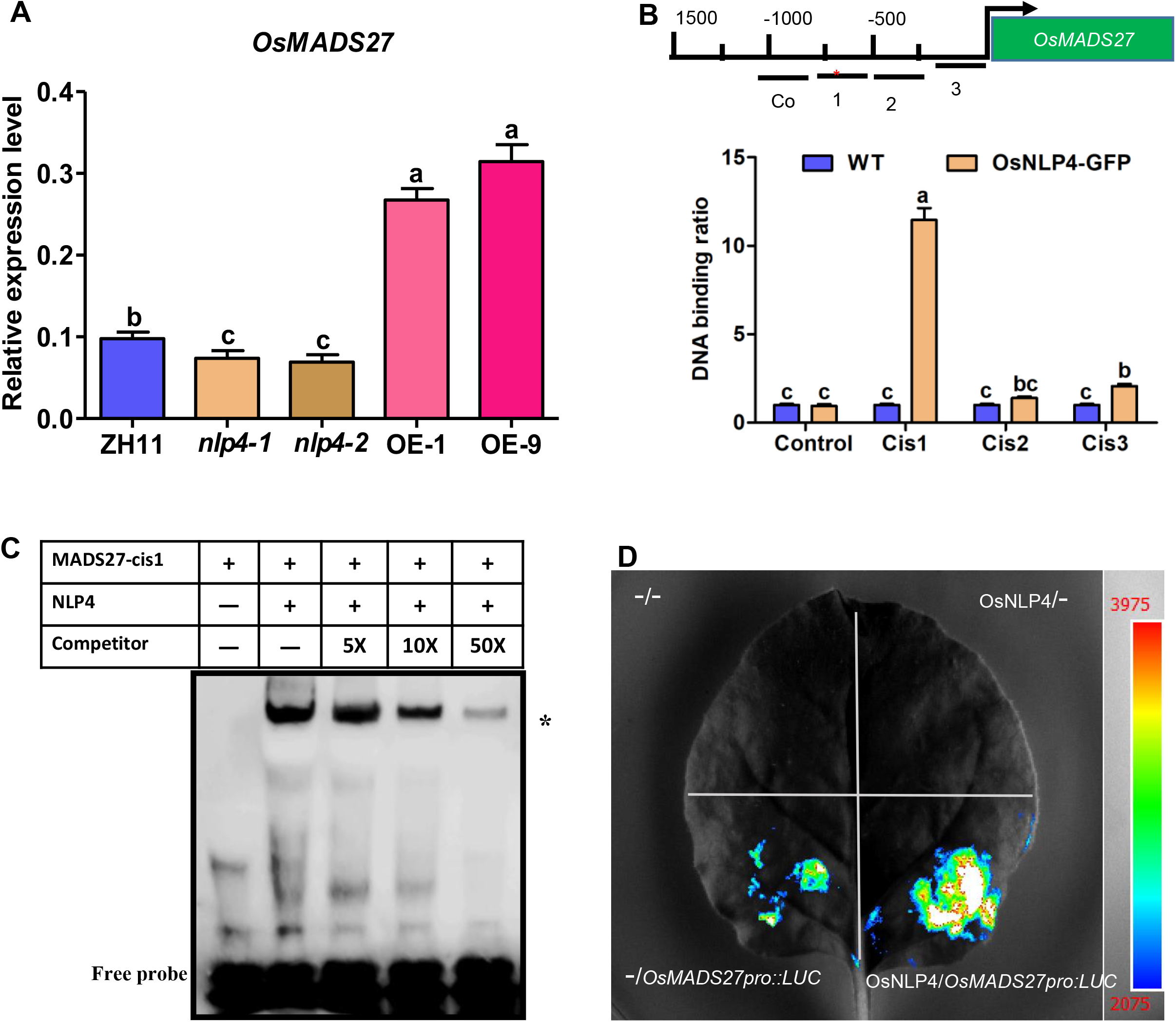
OsNLP4 binds to the promoter of *OsMADS27* and activates its expression. A. qRT-PCR analysis of *OsMADS27* expression in wild type, *nlp4* mutants and *OsNLP4*-OE lines. Seeds were germinated and grown in the hydroponic medium for 16 days before RNA isolation from the whole seedling. Values are mean ± SD (n=3 replicates). Different letters denote significant differences (P < 0.05) from Duncan’s multiple range tests. B. ChIP-qPCR assay. The enrichment of the fragments containing NRE motif (marked with asterisks) in promoters of *OsMADS27* was checked in *OsACTIN1pro:OsNLP4-GFP* plants and wild type. About 200 bp *cis*1 and *cis*3 of *OsMADS27* promoter were enriched in *OsACTIN1pro:OsNLP4-GFP* plants by anti-GFP antibodies as shown in qRT-PCR analyses. Values are mean ± SD (n=3 replicates). Different letters denote significant differences (P < 0.05) from Duncan’s multiple range tests. C. EMSA assay. Recombinant MBP-NLP4 protein was purified from *E. coli* cells and used for DNA binding assays with the promoter of *OsMADS27.* Competition for OsNLP4 binding was conducted with 5×, 10×, 50× unlabeled *OsMADS27* probes. Shifted bands are indicated by asterisk. D. Transcient transactivation assay. pRI101-*OsNLP4* acts as effector. pGreenII0800-*OsMADS27pro::LUC* function as reporters. “-/-” represents pRI101 and pGreenII 0800 empty plasmids. “-/-”, “-/*OsMADS27pro::LUC*”, “OsNLP4/-” as negative controls; “OsNLP4/*OsMADS27pro::LUC*” as experimental group. Different constructs were separately coinfiltrated into 4-week-old tobacco leaves, then the luciferase activity was detected by the luciferase assay system.

**Figure 7.**
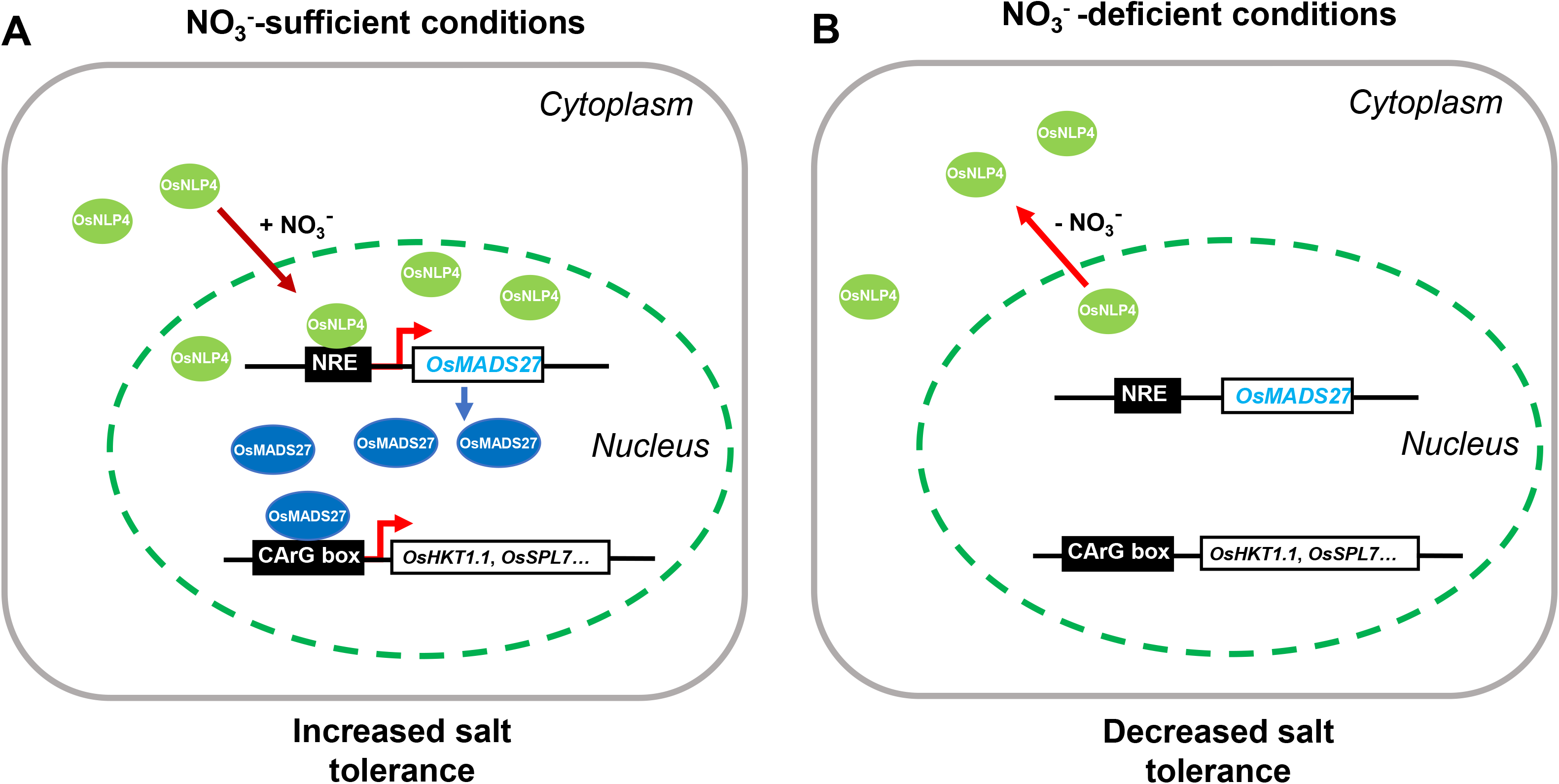
A working model of nitrate-dependent salt tolerance mediated by OsNLP4-OsMADS27 module. A. Under nitrate sufficient condition, nitrate triggers OsNLP4 production and nuclear localization, consequently activating the expression of *OsMADS27*, leading to high level of OsMADS27 that directly binds to the promoters of its target genes such as *OsHKT1.1* and *OsSPL7*, significantly enhancing their expression and improving the salt tolerance of rice. B. Under nitrate deficient condition, less OsNLP4 protein is produced and the vast majority of OsNLP4 protein is localized in the cytoplasm, resulting in a relatively low expression of *OsMADS27*, thereby attenuating the downstream salt tolerance-related genes.

## Discussion

In addition to being an essential nutrient, NO_3_^-^ acts as a signaling molecule involved in controlling multiple metabolic processes in plants (Crawford, 1995). Importantly, nitrate is also a major factor affecting the salt tolerance of crops. NO_3_^-^ application can promote the growth and yield of rice, wheat, canola, citrus, strawberry, pepper, allium, and other plants under salt stress (Çavuşoğlu et al., 2017; Domingo et al., 2004; Gao et al., 2016; Kaya et al., 2003; Kaya and Higgs, 2003; Zheng et al., 2008). However, the intrinsic molecular mechanism of NO_3_^-^-mediated alleviation of salt stress has not been reported so far. In this study, we unraveled the OsNLP4-OsMADS27 module that is crucial for coupling NO_3_^-^ signaling and salt tolerance in rice. We demonstrated that NO_3_^-^ not only induced the expression of *OsMADS27* as described previously (Chen et al., 2018a; Pachamuthu et al., 2022; Puig et al., 2013; Yu et al., 2014), but also promoted the nuclear localization of OsMADS27 (Figs. 1 and 2). OsNLP4, a NO_3_^-^-responsive TF translocating into the nucleus in the presence of NO_3_^-^ (Wu et al., 2021), transcriptionally activates the expression of *OsMADS27* (Fig. 6). Then OsMADS27 activates an array of stress tolerance-related genes as revealed by RNA-seq analyses (Fig. 4) by directly binding to their promoters as demonstrated for *OsHKT1.1* and *OsSPL7* (Fig. 5), thereby enhancing growth and grain yield under salt stress in rice (Figs. 3 and S4). However, in the absence of NO_3_^-^, OsNLP4 was mainly localized in the cytoplasm, resulting in very low expression of *OsMADS27*, which was insufficient to confer salt tolerance in rice as illustrated in the working model (Fig. 8). Our study revealed a novel mechanism of NO_3_^-^ -dependent salt tolerance-mediated by *OsMADS27*, which may be exploited for the improvement of salt tolerance and grain yield in rice.

### Mechanisms of *OsMADS27*-conferred salt tolerance

TFs regulate the expression of various stress-related genes by binding with regulatory motifs in the promoters of these genes in response to stresses (Yamaguchi-Shinozaki and Shinozaki, 2006). Likely benefiting from the simultaneously coordinating the expression of salt-responsive genes (Fig. 4), MADS-box TF *OsMADS27* overexpression increased the transcriptional levels of regulators such as ethylene response factor *OsWR2* (Zhou et al., 2013), salt stress response MYB transcription factor *OsMPS* (Schmidt et al., 2013), A-type response regulator *OsRR2* (Ito and Kurata, 2006), and rice cyclin gene *OsCycB1;3* (La et al., 2006), resulting in significantly improved salt tolerance in germination, seedlings, and reproductive phase of rice (Figs. 3, S3 and S4).

Salt tolerance is highly dependent on intracellular ion homeostasis in order to maintain the turgidity of cell and membrane potential (Bargmann et al., 2009). In our transcriptomic data, the expression of K^+^ transporters such as *OsHKT1.1* (Imran et al., 2020), *OsHKT2.3* (Zhang et al., 2018), K^+^ channel *OsKAT3* (Hwang et al., 2013), and Ca^2+^ sensor *OsCAM1.1* that positively regulates salt tolerance in rice (Saeng-ngam et al., 2012) was significantly enhanced in the OE plants compared with WT under salt stress (Fig. 4C). We found that OsMADS27 directly binds and transcriptionally activates *OsHKT1.1*, which encodes a membrane-localized high-affinity K^+^ transporter (Fig. 5). The *oshkt1.1* knockout mutant rice plants are salt-sensitive depicting its function in the Na^+^ retrieval from leaf blades (Wang et al., 2015). These results demonstrate that *OsMADS27* positively regulates salt tolerance in rice via maintaining ion homeostasis.

In addition, salinity leads to the accumulation of reactive oxygen species (ROS) in plants (Luo et al., 2021), the increased production of which leads to oxidative burden hence being a havoc to cellular membranes as well as macromolecules (Lin et al., 2020). As a target gene of OsMADS27 (Fig. 5), the heat shock transcription factor *OsSPL7* plays an important role in maintaining ROS homeostasis in rice. The *spl7* mutant lost regulation of nicotinamide adenine dinucleotide oxidase, resulting in the accumulation of more H_2_O_2_ in the cells (Hoang et al., 2019). Consistently, our *OsMADS27* overexpression plants exhibited improved resistance against oxidative burden as depicted by our RNA-seq results (Fig. 4C). The upregulation of a number of peroxidases (*OsPRX29*, *OsPRX27*, *OsPRX74*, *OsGPX*, and *OsPRX132*) in OE plants demonstrated that the overexpression of *OsMADS27* ameliorated salt-generated oxidative stress.

ABA, as a stress hormone, plays an important role in the response of plants to salt (Duan et al., 2013; Suzuki et al., 2016b). The enrichment of genes involved in ABA synthesis such as *OsAAO2* and *OsNCED1* (Huang et al., 2021) and ABA-responsive genes such as *OsABI5* and *OsRAB16* (Jiang et al., 2019; Zou et al., 2008) in the OE vs WT group under salt stress (Fig. 4C) implied the possibility that *OsMADS27* may also be involved in ABA signaling. *OsMADS27* has been reported to control NO_3_^-^ -dependent root growth via ABA pathway (Chen et al., 2018a). The possible crosstalk between *OsMADS27*, ABA signaling, and salt stress tolerance needs future attention. Taken together, the salt tolerance mediated by *OsMADS27* in rice is mainly by regulating stress-responsive regulators, balancing ion homeostasis, enhancing ROS scavenging ability, and involving in ABA signaling pathway.

### OsNLP4-OsMADS27 module controls the nitrate dependence of *OsMADS27*-mediated salt tolerance

It is known that multiple members of the MADS-box TF family are involved in the regulation of NO_3_^-^ responses. *Arabidopsis* nitrate regulated1 (AtANR1) is the first NO_3_^-^ regulator found to be involved in the regulation of lateral root developmental plasticity in response to NO_3_^-^ (Zhang and Forde, 1998). *ANR1*-like gene *OsMADS25* is a positive regulator controlling the development of primary and lateral roots of rice by affecting NO_3_^-^ accumulation (Yu et al., 2015). *OsMADS27* is preferentially expressed in roots, and NO_3_^-^ could significantly induce its expression (Yu et al., 2014). We also found that *OsMADS27* specifically responded to NO_3_^-^ rather than ammonium (Fig. 1A, B and 2). The specific NO_3_^-^ responsiveness of *OsMADS27* suggests that a likely NO_3_^-^-responding upstream regulator modulates *OsMADS27*. Indeed, we found that the early NO_3_^-^ response TF OsNLP4 directly binds to the *OsMADS27* promoter and upregulates its expression (Fig. 6). OsNLP4 is a key TF for NO_3_^-^ signaling through nuclear retention mechanisms. Under NO_3_^-^ starvation, OsNLP4 proteins are almost exclusively localized in the cytosol (Wu et al., 2021), hence unable to activate the transcription of *OsMADS27*. However, after NO_3_^-^ was resupplied, OsNLP4 proteins were quickly and predominantly accumulated in the nucleus, resulting in a strong activation of *OsMADS27* (Fig. 1A, B and 6). This OsNLP4-OsMADS27 regulatory module promptly controls *OsMADS27*-mediated salt tolerance in a NO_3_^-^-dependent manner. Recently, it was reported that NO_3_^-^ restriction increased the abundance of miR444, thereby inhibiting the expression of *OsMADS27* and thus regulating rice root development (Pachamuthu et al., 2022). These results indicate that there are multiple ways for NO_3_^-^ signaling to regulate *OsMADS27* expression.

### *OsMADS27* is a positive regulator of grain yield

N uptake and assimilation is closely related to crop yield (Chen et al., 2020; Daniel-Vedele et al., 1998; Hu et al., 2015; Makino, 2011). In addition to controlling salt tolerance in rice, OsMADS27 may also positively regulate grain yield by modulating N metabolism and utilization. In our transcriptomic data, a number of NO_3_^-^ transporters were upregulated in OE compared to WT under salt stress conditions, such as dual affinity NO_3_^-^ transporter OsNRT2.4 (Wei et al., 2018), *OsNAR2.1* required by some members of NRT2 family for NO_3_^-^ transport (Chen et al., 2020), *OsNP5.16,* a positive regulator of grain yield and tiller number (Wang et al., 2021), and low-affinity NO_3_^-^ transporter *OsPTR2* (Li et al., 2015) (Fig. 4C). The significant upregulation of these N transporters and helper protein *OsNAR2.1* correlates with improved yield in transgenic plants under variable N conditions (Fig. S5), suggesting that *OsMADS27* is a positive regulator of rice grain yield.

In conclusion, OsNLP4-OsMADS27 module positively regulates the salt tolerance in rice in a NO_3_^-^ -dependent manner by controlling salt-responsive genes, balancing ion homeostasis, and enhancing ROS scavenging. *OsMADS27* is also an important determinant of yield in rice by modulating the expression of N uptake and assimilation-related genes. Hence, our study fills the gap in the molecular mechanism of NO_3_^-^ -dependent salt tolerance and provides a promising candidate for the development of salt-tolerant crops.

## Methods

### Plant material and culture conditions

The loss-of-function mutants KO1 and KO2 with ZH11 background were generated by Hangzhou Biogle Co., Ltd (Hangzhou, China) (http://www.biogle.cn/), using CRISPR-CAS9 technology, according to the protocol previously described (Lu et al., 2017). The mutants were selected on the bases of their corresponding resistance to hygromycin B. The *ACTIN1*:OsMADS27 overexpression construct was made by inserting the coding region of *OsMADS27* into pCB2006 via GATEWAY cloning system (Lei et al., 2007). The binary vector was transferred into *Agrobacterium tumefaciens* (EHA105) for rice transformation. Homozygous lines (T_3_ generation) were selected using glufosinate and expression was confirmed by RT-PCR and quantitative RT-PCR. These homozygous lines were propagated for obtaining T_4_ generation which was used for further experimental analyses.

A modified Kimura B solution was used for hydroponic culture of rice seedlings in the growth chamber with controlled climate as described (Wu et al., 2021). Growth conditions were maintained at 28°C temperature, photo-regime of 16 hours light /8 hours dark, 70% relative humidity, and light intensity at 250 mmol m^-2^ s^-1^.

### Salt tolerance assays

#### Seed germination

Seeds of wild type, KO1 and KO2 mutants, OE7, and OE8 were washed with distilled water and incubated at 37°C for 7 days. To analyze seed germination, 60∼80 seeds (three replicates per genotype) were randomly placed in a petri dishes containing either water or water plus 150 mM NaCl. The seeds were considered to have germinated when their radicle or germ length reached approximately 1 mm. Seed germination was observed daily to calculate the germination percentage.

#### Seedlings in hydroponic culture

Seeds of wild type, KO1, KO2, OE7, and OE8 were washed with distilled water and incubated at 37°C for 3 days. Germinated seeds were transferred to Hoagland solutions (pH6.0) with different N concentrations (0.02 mM, 0.2 mM, 2 mM KNO_3_) to grow for 7 days, followed by addition of 140 mM NaCl to the culture medium and treated for 7 days. The provided growth conditions were kept at 14-h-light/10-h-dark cycle at 28°C.

#### Seedlings in soil

For the salt treatment in soil, 30 seedlings from each of the wild type, KO1, KO2, OE7, and OE8 were directly grown on soil pot (the pot dimensions were 5×5×12 cm^3^, and five plants were grown per pot). After grown for 4 weeks in soil under greenhouse conditions of 16 h light/ 8 h dark at 30°C, plants were either irrigated with 0 mM or 150 mM NaCl solution for 6-8 days before seedling survival rate was counted.

#### Long-term salt treatment

Seeds of WT, KO1, and OE7 were germinated in plates for 4 days and then transferred to similar pots as previously used for salt treatment of 4-week-old seedlings. The plants were grown in the pots filled with vermiculite and fed with different N concentrations (1.5 mM, 2.5 mM, 5 mM KNO_3_) for 3 weeks, then followed by 65 mM NaCl as salt treatment or without NaCl as control for about 10-12 weeks. Every treatment contains 8 trays with two pots for each genotype and every pot has a single plant. The plants were grown to mature under the greenhouse conditions and yield data were collected.

### RNA extraction and quantitative real-time PCR (qRT-PCR)

Total cellular RNA was extracted from rice tissues (0.08-0.1 g) via Trizol method (Invitrogen, Carlsbad, USA), 1 μg of which was used for cDNA synthesis. The synthesized cDNA was used for qRT-PCR using TaKaRa SYBR Pre-mix Ex-TaqII kit reagents. The primers used are listed in the Table S1. At least three biological replicates were used for each experiment.

### GUS analyses

A 2.0-kb promoter region of *OsMADS27* was amplified from rice genomic DNA (ZH11) followed by its cloning in pCB308R (Lei et al., 2007; Xiang et al., 1999), then recombinant *OsMADS27promotor:GUS* vector was transformed into ZH11 to generate the *OsMADS27:GUS* transgenic plants. For the purpose of GUS staining, the seedlings of *OsMADS27pro:GUS* transgenic plants were incubated in staining solution for 3 hours at 37°C and dehydrated in a series of ethanol (70, 80, 90, and 100%). The dyed tissues were monitored under HiROX MX5040RZ digital optical microscope (Quester China Limited) and then photographed by Nikon D700 digital camera.

### Subcellular localization analyses

The fusion vector *OsMADS27pro*:*OsMADS27-GFP* were created by cloning 2.0-kb promoter and the full length coding sequence of OsMADS27 in the binary vector pUC19. The gene insertion was confirmed by nucleotide sequencing and the ultimate vector was transformed into *Agrobacterium tumefaciens* (EH105). The rice callus was transformed by *Agrobacterium*-based transformation and the selection of positive seedlings was performed by culturing them in hygromycin B containing medium. To investigate the nuclear-cytoplasmic shuttling of OsMADS27, the positive seedlings were grown on the modified Kimura B solution with 2 mM KNO_3_ or without N for 10 days. Subsequently, N-free medium treated with either 2 mM KNO_3_ or 2 mM NH_4_Cl or 150 mM NaCl for 60 min, and returned to the N-free medium. Additionally, 2 mM KNO_3_ medium treated with 150 mM NaCl for 60 min, and returned to N-free medium. The confocal microscopy was performed by Zeiss 710 microscope having argon laser (488 nm for GFP excitation).

### Western blot analysis

Proteins were extracted from the 2-week-old seedlings grown hydroponically on medium containing different N concentrations 0.02 mM, 0.2 mM and 2 mM KNO_3_ without salt as control or with 100 mM NaCl using the RIPA lysis buffer (strong) (Beyotime, China). For western blot analysis, proteins were electroblotted from 10% acrylamide gel to nitrocellulose membrane (Immobilon-P, MILLIPORE Corporation, Bedford, MA, USA) after the separation of SDS-PAGE. Antibodies used in western blot were as follows: anti-GFP antibody (M20004, Mouse mAb, Abmart, Shanghai, China), 1:1000 for western blot; anti-ACTIN antibody (M20009, Mouse mAb, Abmart, Shanghai, China), 1:1000 for western blot and goat anti-mouse lgG-HRP (M21001, Abmart, Shanghai, China), 1:5000 for western blot. Image Quant LAS 4000 (GE, USA), as the CCD camera system, was used for the band intensity quantification with Super Signal West Femto Trial Kit (Thermo, Rockford, IL, USA).

### Electrophoretic mobility shift assay (EMSA)

Electrophoretic mobility shift assay (EMSA) was conducted as previously described (Hellman and Fried, 2007). The coding sequence of *OsNLP4* was cloned into pMAL c2x vector and MBP-NLP4 fusion protein was expressed in *E. coli* Rosseta2 strain. Biotin-labelled DNA that 45 bp fragment containing CArG motif and unlabelled competitor DNA were synthesized by Sangon Biotech Co., Ltd (Shang Hai, China). DNA probes were generated by cooling down the mixtures of complementary oligonucleotides from 95°C to room temperature. EMSA assay was performed using a LightShift™ EMSA Optimization and Control Kit (20148×) (Thermo Fisher Scientific, Waltham, USA). The reaction mixtures were loaded on 6% polyacrylamide gel in 0.5 × TBE buffer and electrophoresed at 4°C. These results were detected by a CCD camera system (Image Quant LAS 4000).

### Transient transactivation assays in tobacco leaf

Transient transactivation assay in tobacco leaf was performed as previously described (Lim et al., 2017). The coding sequences of *OsMADS27/OsNLP4* were constructed into pRI101 vector as reporters. About 2500 bp promoters of *OsHKT1.1*, *OsSPL7*, *OsMADS27* were respectively cloned into pGreenII 0800 vector as reporters. These constructs were electroporated into *Agrobacterium* GV3101 strain, then cultured in LB medium at 28°C for 2 days. The precipitate was collected by centrifugation at 5000 rpm for 5 min, resuspended with infiltration buffer (10 mM MES, 10 mM MgCl_2_, 150 mM acetosyringone, pH 5.6), and incubated at room temperature for 2 hours before co-injecting into *Nicotiana benthamiana* leaves. 3 days after injection, tobacco leaves were sprayed using LUC substrates (1 mM Xeno lightTM D-luciferin potassium salt). At least three biological replicates were used for each experiment.

### RNA-sequencing analysis

Each genotype has about 100 seedlings (ZH11 background) of every treatment was grown hydroponically in a growth chamber with the condition described above. The seedlings were cultured in modified Kimura B solution with 1.5 mM KNO_3_ for 12 days and treated with 100 mM NaCl or without NaCl as a control for another three days. 15-day-old seedlings (whole plants) were sampled for RNA-sequencing. For each treatment, 20 seedlings were collected as a sample, and three independent biological replicates were conducted. RNA library construction and sequence analysis were conducted as described previously (Khan et al., 2016a).

### Yeast one-hybrid assay

The protein putative binding sites were cloned into BD vector (pHIS2), and the coding sequences of *OsMADS27/OsNLP4* were cloned into AD vector (pAD-GAL4-2.1), respectively. A yeast one-hybrid (Y1H) assay was conducted according to the procedure described previously (Mao et al., 2016).

### Chromatin immunoprecipitation–quantitative PCR assay

A chromatin immunoprecipitation (ChIP) assay was carried out according to the protocol described before (O’Geen et al., 2010) with minor modifications. Transgenic rice (*OsMADS27pro:OsMADS27-GFP*) seedlings were grown under high nitrogen (2 mM) condition for 2 weeks. About 2.0 g of seedlings were placed in 1% formaldehyde (v/v) at 20-25 °C in vacuum for 15 min and then homogenized within liquid nitrogen. Chromatin from lysed nuclei was fragmented ultrasonically to achieve an average length of 500 bp. The anti-GFP antibodies (Sigma, F1804) were immunoprecipitated overnight at 4 °C. The immuno-precipitated DNA fragments were dissolved in water and kept at −80 °C before use. The precipitated fragments were used as template for quantitative PCR (qPCR).

### Field trial of rice

For the field test of KO1 mutant and *OsMADS27*-overexpressing (OE7) (all with ZH11 background), T3 generation plants were grown in Chang Xing, Zhejiang in 2021 (April to September). The plant density was 6 rows. 20 plants per row for each plot, and four replicates were used for each N condition. Urea was used as the N fertilizer at 94 kg N hm^−2^ for low N (LN), 184 kg N hm^−2^ for normal N (NN), and 375 kg N hm^−2^ for high N (HN). To reduce the variability in the field test, the fertilizers were used evenly in each plot for N application level. The plants at the edge were excluded from data collection in each plot in order to avoid margin effects.

### Agronomic trait analyses

Individual tiller number, panicle number, and grain yield per plant were measured according to a protocol documented earlier (Hu et al., 2015).

### Accession numbers

Sequence data from this article can be found in the Rice Genome Annotation Project (https://rice.plantbiology.msu.edu/) under the following accession numbers: *OsMADS27*, *LOC_Os02g36924*; *OsHKT1.1*, *LOC_Os04g51820*; *OsNLP4*, *LOC_Os09g37710*; *OsSPL7*, *LOC_Os05g45410*; *OsHKT2.3*, *LOC_Os01g34850*; *OsKAT3*, *LOC_Os02g14840*; *OsO3L2*, *LOC_Os06g36390*; *OsMPS*, *LOC_Os02g40530*.

## Supplemental Data

Fig. S1. Expression pattern of *OsMADS27*.

Fig. S2. Verification of the CRISPR/Cas9-edited mutations in *OsMADS27* and overexpression lines of *OsMADS27*.

Fig. S3. *OsMADS27* positively affects salt tolerance in germination and seedling growth.

Fig. S4. OsMADS7 positively affects grain yield in a N-dependent manner under normal and salt stress conditions.

Fig. S5. *OsMADS27* improves NUE and grain yield in the field of different nitrogen concentrations.

Fig. S6. The gene ontology (GO)-based enrichment analysis of DEGs.

Fig. S7. *OsMADS27* broadly regulates the genes involvedin salt tolerance.

Table S1. List of primers used in this study.

## Author Contributions

C.B.X., A.A., P.X.Z., and J.W. designed the experiments. A.A., T.N., J.Z., J.W., Y.S., P.X.Z. and S.U.J. performed experiments and data analyses. J.W., Z.S.Z., J.Q.X. and Z.Y.Z. performed field trials and data analyses. A.A. and J.W. wrote the manuscript. C.B.X., P.X.Z. and J.W. revised the manuscript. C.B.X. supervised the project.

## Acknowledgements

This work was supported by the Strategic Priority Research Program of the Chinese Academy of Sciences (grant no. XDA24010303). Alamin Alfatih was a recipient of CAS-TWAS President’s Fellowship and CAS International Postdoctoral Fellowship. Sami Ullah Jan was a recipient of CAS-TWAS President’s Fellowship. The authors declare no conflicts of interest.

## References

Adem, G.D., Roy, S.J., Zhou, M., Bowman, J.P. and Shabala, S. (2014) Evaluating contribution of ionic, osmotic and oxidative stress components towards salinity tolerance in barley. BMC plant biology 14, 113.

Ahammed, G.J., Li, X., Yang, Y., Liu, C., Zhou, G., Wan, H. and Cheng, Y. (2020) Tomato WRKY81 acts as a negative regulator for drought tolerance by modulating guard cell H2O2–mediated stomatal closure. Environmental and Experimental Botany 171, 103960.

Aragao, R.M., Silva, E.N., Vieira, C.F. and Silveira, J.A. (2012) High supply of NO 3− mitigates salinity effects through an enhancement in the efficiency of photosystem II and CO 2 assimilation in Jatropha curcas plants. Acta Physiologiae Plantarum 34, 2135–2143.

Asano, T., Hayashi, N., Kobayashi, M., Aoki, N., Miyao, A., Mitsuhara, I., Ichikawa, H., Komatsu, S., Hirochika, H., Kikuchi, S. and Ohsugi, R. (2012) A rice calcium-dependent protein kinase OsCPK12 oppositely modulates salt-stress tolerance and blast disease resistance. Plant Journal 69, 26–36.

Ashraf, M., Athar, H., Harris, P. and Kwon, T. (2008) Some prospective strategies for improving crop salt tolerance. Advances in agronomy 97, 45–110.

Bargmann, B.O., Laxalt, A.M., Riet, B.t., Van Schooten, B., Merquiol, E., Testerink, C., Haring, M.A., Bartels, D. and Munnik, T. (2009) Multiple PLDs required for high salinity and water deficit tolerance in plants. Plant and Cell Physiology 50, 78–89.

Bose, J., Rodrigo-Moreno, A. and Shabala, S. (2014) ROS homeostasis in halophytes in the context of salinity stress tolerance. Journal of experimental botany 65, 1241–1257.

Campo, S., Baldrich, P., Messeguer, J., Lalanne, E., Coca, M. and San Segundo, B. (2014) Overexpression of a Calcium-Dependent Protein Kinase Confers Salt and Drought Tolerance in Rice by Preventing Membrane Lipid Peroxidation. Plant Physiology 165, 688–704.

Cao, M.J., Wang, Z., Zhao, Q., Mao, J.L., Speiser, A., Wirtz, M., Hell, R., Zhu, J.K. and Xiang, C.B. (2014) Sulfate availability affects ABA levels and germination response to ABA and salt stress in Arabidopsis thaliana. Plant Journal 77, 604–615.

Çavuşoğlu, K., Cadıl, S. and Çavuşoğlu, D. (2017) Role of Potassium Nitrate (KNO_3_) in Alleviation of Detrimental Effects of Salt Stress on Some Physiological and Cytogenetical Parameters in Allium cepa L. Cytologia 82, 279–286.

Chakraborty, K., Bose, J., Shabala, L. and Shabala, S. (2016) Difference in root K+ retention ability and reduced sensitivity of K+-permeable channels to reactive oxygen species confer differential salt tolerance in three Brassica species. Journal of Experimental Botany 67, 4611–4625.

Chen, C., Begcy, K., Liu, K., Folsom, J.J., Wang, Z., Zhang, C. and Walia, H. (2016) Heat stress yields a unique MADS box transcription factor in determining seed size and thermal sensitivity. Plant physiology 171, 606–622.

Chen, G., Hu, Q., Luo, L., Yang, T., Zhang, S., Hu, Y., Yu, L. and Xu, G. (2015) Rice potassium transporter OsHAK1 is essential for maintaining potassium-mediated growth and functions in salt tolerance over low and high potassium concentration ranges. Plant Cell and Environment 38, 2747–2765.

Chen, H., Xu, N., Wu, Q., Yu, B., Chu, Y., Li, X., Huang, J. and Jin, L. (2018a) OsMADS27 regulates the root development in a NO3−—Dependent manner and modulates the salt tolerance in rice (Oryza sativa L.). Plant Science 277, 20–32.

Chen, J., Liu, X., Liu, S., Fan, X., Zhao, L., Song, M., Fan, X. and Xu, G. (2020) Co-overexpression of OsNAR2. 1 and OsNRT2. 3a increased agronomic nitrogen use efficiency in transgenic rice plants. Frontiers in Plant Science 11, 1245.

Chen, L., Zhao, Y., Xu, S., Zhang, Z., Xu, Y., Zhang, J. and Chong, K. (2018b) Os MADS 57 together with Os TB 1 coordinates transcription of its target Os WRKY 94 and D14 to switch its organogenesis to defense for cold adaptation in rice. New Phytologist 218, 219–231.

Chen, Z., Zhao, P.X., Miao, Z.Q., Qi, G.F., Wang, Z., Yuan, Y., Ahmad, N., Cao, M.J., Hell, R., Wirtz, M. and Xiang, C.B. (2019) SULTR3s Function in Chloroplast Sulfate Uptake and Affect ABA Biosynthesis and the Stress Response. Plant Physiology 180, 593–604.

Clarkson, D.T. and Hanson, J.B. (1980) The mineral nutrition of higher plants. Annual Review of Plant Biology 31, 239–298.

Crawford, N.M. (1995) Nitrate: Nutrient and Signal for Plant Growth. Plant Cell 7, 859–868.

Daniel-Vedele, F., Filleur, S. and Caboche, M. (1998) Nitrate transport: a key step in nitrate assimilation. Current Opinion in Plant Biology 1, 235–239.

Domingo, J.I., Yoseph, L., Aurelio, G.C., Francisco, R.T., Eduardo, P.M. and Manuel, T. (2004) Nitrate improves growth in salt-stressed citrus seedlings through effects on photosynthetic activity and chloride accumulation. Tree Physiology 24, 1027–1034.

Duan, L., Dietrich, D., Ng, C.H., Chan, P.M., Bhalerao, R., Bennett, M.J. and Dinneny, J.R. (2013) Endodermal ABA signaling promotes lateral root quiescence during salt stress in Arabidopsis seedlings. Plant Cell 25, 324–341.

Fatma, M., Asgher, M., Masood, A. and Khan, N.A. (2014) Excess sulfur supplementation improves photosynthesis and growth in mustard under salt stress through increased production of glutathione. Environmental and Experimental Botany 107, 55–63.

Fatma, M., Iqbal, N., Gautam, H., Sehar, Z., Sofo, A., D’Ippolito, I. and Khan, N.A. (2021) Ethylene and Sulfur Coordinately Modulate the Antioxidant System and ABA Accumulation in Mustard Plants under Salt Stress. Plants 10, 180.

Gao, L., Liu, M., Wang, M., Shen, Q. and Guo, S. (2016) Enhanced Salt Tolerance under Nitrate Nutrition is Associated with Apoplast Na+ Content in Canola (Brassica. napus L.) and Rice (Oryza sativa L.) Plants. Plant Cell Physiology 57, 2323–2333.

Guo, J., Zhou, Q., Li, X., Yu, B. and Luo, Q. (2017) Enhancing NO3-supply confers NaCl tolerance by adjusting Cl-uptake and transport in G. max & G. soja. Journal of soil science and plant nutrition 17, 194–202.

Hamamoto, S., Horie, T., Hauser, F., Deinlein, U., Schroeder, J.I. and Uozumi, N. (2015) HKT transporters mediate salt stress resistance in plants: from structure and function to the field. Current opinion in biotechnology 32, 113–120.

Hellman, L.M. and Fried, M.G. (2007) Electrophoretic mobility shift assay (EMSA) for detecting protein-nucleic acid interactions. Nat Protoc 2, 1849–1861.

Hoang, T.V., Vo, K.T.X., Rahman, M.M., Choi, S.-H. and Jeon, J.-S. (2019) Heat stress transcription factor OsSPL7 plays a critical role in reactive oxygen species balance and stress responses in rice. Plant Science 289, 110273.

Hu, B., Wang, W., Ou, S., Tang, J., Li, H., Che, R., Zhang, Z., Chai, X., Wang, H. and Wang, Y. (2015) Variation in NRT1. 1B contributes to nitrate-use divergence between rice subspecies. Nature Genetics 47, 834-838.

Huang, S., Liang, Z., Chen, S., Sun, H., Fan, X., Wang, C., Xu, G. and Zhang, Y. (2019) A transcription factor, OsMADS57, regulates long-distance nitrate transport and root elongation. Plant Physiology 180, 882-895.

Huang, Y.t., Wu, W., Zhao, T.y., Lu, M., Wu, H.p. and Cao, D.d. (2021) Drying temperature regulates vigor of high moisture rice seeds via involvement in phytohormone, ROS, and relevant gene expression. Journal of the Science of Food and Agriculture 101, 2143–2155.

Hwang, H., Yoon, J., Kim, H.Y., Min, M.K., Kim, J.-A., Choi, E.-H., Lan, W., Bae, Y.-M., Luan, S. and Cho, H. (2013) Unique features of two potassium channels, OsKAT2 and OsKAT3, expressed in rice guard cells. PLoS One 8, e72541.

Imran, S., Horie, T. and Katsuhara, M. (2020) Expression and ion transport activity of rice OsHKT1; 1 variants. Plants 9, 16.

Iqbal, N., Umar, S. and Khan, N.A. (2015) Nitrogen availability regulates proline and ethylene production and alleviates salinity stress in mustard (Brassica juncea). Journal of Plant Physiology 178, 84–91.

Ito, Y. and Kurata, N. (2006) Identification and characterization of cytokinin-signalling gene families in rice. Gene 382, 57–65.

Jiang, D., Zhou, L., Chen, W., Ye, N., Xia, J. and Zhuang, C. (2019) Overexpression of a microRNA-targeted NAC transcription factor improves drought and salt tolerance in Rice via ABA-mediated pathways. Rice 12, 1–11.

Kaya, C., Ak, B.E. and Higgs, D. (2003) Response of Salt-Stressed Strawberry Plants to Supplementary Calcium Nitrate and/or Potassium Nitrate. Journal of Plant Nutrition 26, 543–560.

Kaya, C. and Higgs, D. (2003) Supplementary Potassium Nitrate Improves Salt Tolerance in Bell Pepper Plants. Journal of Plant Nutrition 26, 1367–1382.

Kaya, C., Tuna, A.L., Ashraf, M. and Altunlu, H. (2007) Improved salt tolerance of melon (Cucumis melo L.) by the addition of proline and potassium nitrate. Environmental and Experimental Botany 60, 397–403.

Khan, A.U.H., Rathore, M.G., Allende-Vega, N., Vo, D.-N., Belkhala, S., Orecchioni, S., Talarico, G., Bertolini, F., Cartron, G. and Lecellier, C.-H. (2016a) Human leukemic cells performing oxidative phosphorylation (OXPHOS) generate an antioxidant response independently of reactive oxygen species (ROS) production. EBioMedicine 3, 43–53.

Khan, H., Ashraf, M., Shahzad, S., Aziz, A., Piracha, M. and Siddiqui, A. (2016b) Adequate regulation of plant nutrients for improving cotton adaptability to salinity stress. J Appl Agric Biotechnol 1, 47–56.

Khong, G.N., Pati, P.K., Richaud, F., Parizot, B., Bidzinski, P., Mai, C.D., Bès, M., Bourrié, I., Meynard, D. and Beeckman, T. (2015) OsMADS26 negatively regulates resistance to pathogens and drought tolerance in rice. Plant Physiology 169, 2935–2949.

Konishi, M. and Yanagisawa, S. (2010) Identification of a nitrate-responsive cis-element in the Arabidopsis NIR1 promoter defines the presence of multiple cis-regulatory elements for nitrogen response. Plant Journal 63, 269–282.

Kumar, K., Kumar, M., Kim, S.-R., Ryu, H. and Cho, Y.-G. (2013) Insights into genomics of salt stress response in rice. Rice 6, 1–15.

La, H., Li, J., Ji, Z., Cheng, Y., Li, X., Jiang, S., Venkatesh, P.N. and Ramachandran, S. (2006) Genome-wide analysis of cyclin family in rice (Oryza Sativa L.). Molecular Genetics and Genomics 275, 374–386.

Lei, Z.Y., Zhao, P., Cao, M.J., Cui, R., Chen, X., Xiong, L.Z., Zhang, Q.F., Oliver, D.J. and Xiang, C.B. (2007) High-throughput binary vectors for plant gene function analysis. Journal of Integrative Plant Biology 49, 556–567.

Li, Y., Ouyang, J., Wang, Y.-Y., Hu, R., Xia, K., Duan, J., Wang, Y., Tsay, Y.-F. and Zhang, M. (2015) Disruption of the rice nitrate transporter OsNPF2. 2 hinders root-to-shoot nitrate transport and vascular development. Scientific reports 5, 1-10.

Lim, S.H., Kim, D.H., Kim, J.K., Lee, J.Y. and Ha, S.H. (2017) A radish basic helix-loop-helix transcription factor, RsTT8 acts a positive regulator for anthocyanin biosynthesis. Front Plant Sci 8, 1917.

Lin, H.X., Yang, Y.Q., Quan, R.D., Mendoza, I., Wu, Y.S., Du, W.M., Zhao, S.S., Schumaker, K.S., Pardo, J.M. and Guo, Y. (2009) Phosphorylation of SOS3-LIKE CALCIUM BINDING PROTEIN8 by SOS2 Protein Kinase Stabilizes Their Protein Complex and Regulates Salt Tolerance in Arabidopsis. Plant Cell 21, 1607–1619.

Lin, Y.-J., Yu, X.-Z., Li, Y.-H. and Yang, L. (2020) Inhibition of the mitochondrial respiratory components (Complex I and Complex III) as stimuli to induce oxidative damage in Oryza sativa L. under thiocyanate exposure. Chemosphere 243, 125472.

Lu, Y., Ye, X., Guo, R., Huang, J., Wang, W., Tang, J., Tan, L., Zhu, J.-k., Chu, C. and Qian, Y. (2017) Genome-wide targeted mutagenesis in rice using the CRISPR/Cas9 system. Molecular Plant 10, 1242–1245.

Luo, X., Dai, Y., Zheng, C., Yang, Y., Chen, W., Wang, Q., Chandrasekaran, U., Du, J., Liu, W. and Shu, K. (2021) The ABI4-RbohD/VTC2 regulatory module promotes reactive oxygen species (ROS) accumulation to decrease seed germination under salinity stress. New Phytologist 229, 950–962.

Makino, A. (2011) Photosynthesis, grain yield, and nitrogen utilization in rice and wheat. Plant Physiol 155, 125–129.

Manishankar, P., Wang, N., Koster, P., Alatar, A.A. and Kudla, J. (2018) Calcium Signaling during Salt Stress and in the Regulation of Ion Homeostasis. Journal of Experimental Botany 69, 4215–4226.

Mansour, M. (2000) Nitrogen containing compounds and adaptation of plants to salinity stress. Biologia Plantarum 43, 491–500.

Mao, J.-L., Miao, Z.-Q., Wang, Z., Yu, L.-H., Cai, X.-T. and Xiang, C.-B. (2016) Arabidopsis ERF1 mediates cross-talk between ethylene and auxin biosynthesis during primary root elongation by regulating ASA1 expression. PLoS genetics 12, e1005760.

Martínez-Atienza, J., Jiang, X., Garciadeblas, B., Mendoza, I., Zhu, J.-K., Pardo, J.M. and Quintero, F.J. (2007) Conservation of the salt overly sensitive pathway in rice. Plant physiology 143, 1001–1012.

Moyle, R., Fairbairn, D.J., Ripi, J., Crowe, M. and Botella, J.R. (2005) Developing pineapple fruit has a small transcriptome dominated by metallothionein. Journal of Experimental Botany 56, 101–112.

Munns, R. and Tester, M. (2008) Mechanisms of salinity tolerance. Annu. Rev. Plant Biol. 59, 651–681.

Nasab, A.R., Pour, A.T. and Shirani, H. (2014) Effect of salinity and nitrogen application on growth, chemical composition and some biochemical indices of pistachio seedlings (Pistacia vera L.). Journal of Plant Nutrition 37, 1612-1626.

O’Geen, H., Frietze, S. and Farnham, P.J. (2010) Using ChIP-seq technology to identify targets of zinc finger transcription factors. In: Engineered Zinc Finger Proteins pp. 437-455. Springer.

Pachamuthu, K., Hari Sundar, V., Narjala, A., Singh, R.R., Das, S., Avik Pal, H.C.Y. and Shivaprasad, P.V. (2022) Nitrate-dependent regulation of miR444-OsMADS27 signaling cascade controls root development in rice. Journal of Experimental Botany erac083.

Puig, J., Meynard, D., Khong, G.N., Pauluzzi, G., Guiderdoni, E. and Gantet, P. (2013) Analysis of the expression of the AGL17-like clade of MADS-box transcription factors in rice. Gene Expression Patterns 13, 160–170.

Qadir, M., Quillérou, E., Nangia, V., Murtaza, G., Singh, M., Thomas, R.J., Drechsel, P. and Noble, A.D. (2014) Economics of salt-induced land degradation and restoration. In: Natural resources forum pp. 282–295. Wiley Online Library.

Qiu, Q.S., Guo, Y., Dietrich, M.A., Schumaker, K.S. and Zhu, J.K. (2002) Regulation of SOS1, a plasma membrane Na+/H+ exchanger in Arabidopsis thaliana, by SOS2 and SOS3. P Natl Acad Sci USA 99, 8436–8441.

Raddatz, N., Morales de Los Rios, L., Lindahl, M., Quintero, F.J. and Pardo, J.M. (2020) Coordinated Transport of Nitrate, Potassium, and Sodium. Frontiers in Plant Science 11, 247.

Rais, L., Masood, A., Inam, A. and Khan, N. (2013) Sulfur and nitrogen co-ordinately improve photosynthetic efficiency, growth and proline accumulation in two cultivars of mustard under salt stress. Journal of Plant Biochemistry & Physiology.

Rosas-Santiago, P., Lagunas-Gómez, D., Barkla, B.J., Vera-Estrella, R., Lalonde, S., Jones, A., Frommer, W.B., Zimmermannova, O., Sychrová, H. and Pantoja, O. (2015) Identification of rice cornichon as a possible cargo receptor for the Golgi-localized sodium transporter OsHKT1; 3. Journal of experimental botany 66, 2733–2748.

Saeng-ngam, S., Takpirom, W., Buaboocha, T. and Chadchawan, S. (2012) The role of the OsCam1-1 salt stress sensor in ABA accumulation and salt tolerance in rice. Journal of Plant Biology 55, 198–208.

Schmidt, R., Schippers, J.H., Mieulet, D., Obata, T., Fernie, A.R., Guiderdoni, E. and Mueller-Roeber, B. (2013) MULTIPASS, a rice R2R3-type MYB transcription factor, regulates adaptive growth by integrating multiple hormonal pathways. Plant Journal 76, 258–273.

Suzuki, K., Yamaji, N., Costa, A., Okuma, E., Kobayashi, N.I., Kashiwagi, T., Katsuhara, M., Wang, C., Tanoi, K. and Murata, Y. (2016a) OsHKT1; 4-mediated Na+ transport in stems contributes to Na+ exclusion from leaf blades of rice at the reproductive growth stage upon salt stress. BMC plant biology 16, 1–15.

Suzuki, N., Bassil, E., Hamilton, J.S., Inupakutika, M.A., Zandalinas, S.I., Tripathy, D., Luo, Y., Dion, E., Fukui, G., Kumazaki, A., Nakano, R., Rivero, R.M., Verbeck, G.F., Azad, R.K., Blumwald, E. and Mittler, R. (2016b) ABA Is Required for Plant Acclimation to a Combination of Salt and Heat Stress. PLoS One 11, e0147625.

Tian, Q., Shen, L., Luan, J., Zhou, Z., Guo, D., Shen, Y., Jing, W., Zhang, B., Zhang, Q. and Zhang, W. (2021) Rice shaker potassium channel OsAKT2 positively regulates salt tolerance and grain yield by mediating K(+) redistribution. Plant Cell and Environment 44, 2951–2965.

Wang, J., Wan, R., Nie, H., Xue, S. and Fang, Z. (2021) OsNPF5. 16, a nitrate transporter gene with natural variation, is essential for rice growth and yield. The Crop Journal.

Wang, R., Jing, W., Xiao, L., Jin, Y., Shen, L. and Zhang, W. (2015) The rice high-affinity potassium transporter1; 1 is involved in salt tolerance and regulated by an MYB-type transcription factor. Plant Physiology 168, 1076–1090.

Wei, J., Zheng, Y., Feng, H., Qu, H., Fan, X., Yamaji, N., Ma, J.F. and Xu, G. (2018) OsNRT2. 4 encodes a dual-affinity nitrate transporter and functions in nitrate-regulated root growth and nitrate distribution in rice. Journal of experimental botany 69, 1095-1107.

Wu, H., Zhang, X., Giraldo, J.P. and Shabala, S. (2018) It is not all about sodium: revealing tissue specificity and signalling roles of potassium in plant responses to salt stress. Plant and Soil 431, 1–17.

Wu, J., Yu, C., Hunag, L., Wu, M., Liu, B., Liu, Y., Song, G., Liu, D. and Gan, Y. (2020) Overexpression of MADS-box transcription factor OsMADS25 enhances salt stress tolerance in Rice and Arabidopsis. Plant Growth Regulation 90, 163–171.

Wu, J., Zhang, Z.S., Xia, J.Q., Alfatih, A., Song, Y., Huang, Y.J., Wan, G.Y., Sun, L.Q., Tang, H., Liu, Y., Wang, S.M., Zhu, Q.S., Qin, P., Wang, Y.P., Li, S.G., Mao, C.Z., Zhang, G.Q., Chu, C., Yu, L.H. and Xiang, C.B. (2021) Rice NIN-LIKE PROTEIN 4 plays a pivotal role in nitrogen use efficiency. Plant Biotechnol Journal 19, 448–461.

Wu, R., Tomes, S., Karunairetnam, S., Tustin, S.D., Hellens, R.P., Allan, A.C., Macknight, R.C. and Varkonyi-Gasic, E. (2017) SVP-like MADS Box Genes Control Dormancy and Budbreak in Apple. Frontiers in Plant Science 8, 477.

Xiang, C., Han, P., Lutziger, I., Wang, K. and Oliver, D.J. (1999) A mini binary vector series for plant transformation. Plant molecular biology 40, 711–717.

Yamaguchi-Shinozaki, K. and Shinozaki, K. (2006) Transcriptional regulatory networks in cellular responses and tolerance to dehydration and cold stresses. Annu. Rev. Plant Biol. 57, 781–803.

Yang, Y.Q. and Guo, Y. (2018a) Elucidating the molecular mechanisms mediating plant salt-stress responses. New Phytol 217, 523–539.

Yang, Y.Q. and Guo, Y. (2018b) Unraveling salt stress signaling in plants. J Integr Plant Biol 60, 796–804.

Yin, X., Liu, X., Xu, B., Lu, P., Dong, T., Yang, D., Ye, T., Feng, Y.Q. and Wu, Y. (2019) OsMADS18, a membrane-bound MADS-box transcription factor, modulates plant architecture and the abscisic acid response in rice. Journal of Experimental Botany 70, 3895-3909.

Yu, C., Liu, Y., Zhang, A., Su, S., Yan, A., Huang, L., Ali, I., Liu, Y., Forde, B.G. and Gan, Y. (2015) MADS-box transcription factor OsMADS25 regulates root development through affection of nitrate accumulation in rice. PLoS One 10, e0135196.

Yu, C., Su, S., Xu, Y., Zhao, Y., Yan, A., Huang, L., Ali, I. and Gan, Y. (2014) The effects of fluctuations in the nutrient supply on the expression of five members of the AGL17 clade of MADS-box genes in rice. PLoS One 9, e105597.

Yu, L.-H., Wu, J., Zhang, Z.-S., Miao, Z.-Q., Zhao, P.-X., Wang, Z. and Xiang, C.-B. (2017) Arabidopsis MADS-box transcription factor AGL21 acts as environmental surveillance of seed germination by regulating ABI5 expression. Molecular plant 10, 834–845.

Zhang, H. and Forde, B.G. (1998) An Arabidopsis MADS box gene that controls nutrient-induced changes in root architecture. Science 279, 407–409.

Zhang, H., Yang, B., Liu, J., Guo, D., Hou, J., Chen, S., Song, B. and Xie, C. (2017) Analysis of structural genes and key transcription factors related to anthocyanin biosynthesis in potato tubers. Scientia Horticulturae 225, 310–316.

Zhang, Y., Fang, J., Wu, X. and Dong, L. (2018) Na+/K+ balance and transport regulatory mechanisms in weedy and cultivated rice (Oryza sativa L.) under salt stress. BMC plant biology 18, 1–14.

Zhao, P.-X., Miao, Z.-Q., Zhang, J., Chen, S.-Y., Liu, Q.-Q. and Xiang, C.-B. (2020) Arabidopsis MADS-box factor AGL16 negatively regulates drought resistance via stomatal density and stomatal movement. Journal of Experimental Botany 71, 6092–6106.

Zhao, P.X., Zhang, J., Chen, S.Y., Wu, J., Xia, J.Q., Sun, L.Q., Ma, S.S. and Xiang, C.B. (2021) Arabidopsis MADS-box factor AGL16 is a negative regulator of plant response to salt stress by downregulating salt-responsive genes. New Phytologist 232, 2418–2439.

Zheng, Y., Jia, A., Ning, T., Xu, J., Li, Z. and Jiang, G. (2008) Potassium nitrate application alleviates sodium chloride stress in winter wheat cultivars differing in salt tolerance. J Plant Physiol 165, 1455–1465.

Zhou, X., Jenks, M.A., Liu, J., Liu, A., Zhang, X., Xiang, J., Zou, J., Peng, Y. and Chen, X. (2013) Overexpression of Transcription Factor OsWR2 Regulates Wax and Cutin Biosynthesis in Rice and Enhances its Tolerance to Water Deficit. Plant Molecular Biology Reporter 32, 719–731.

Zhu, J.K., Liu, J.P. and Xiong, L.M. (1998) Genetic analysis of salt tolerance in Arabidopsis: Evidence for a critical role of potassium nutrition. Plant Cell 10, 1181–1191.

Zorb, C., Senbayram, M. and Peiter, E. (2014) Potassium in Agriculture: Status and Perspectives. Journal of Plant Physiology 171, 656–669.

Zou, M., Guan, Y., Ren, H., Zhang, F. and Chen, F. (2008) A bZIP transcription factor, OsABI5, is involved in rice fertility and stress tolerance. Plant molecular biology 66, 675-683.

