## supplemental info for "Nitrate-dependent salt tolerance mediated by OsNLP4-OsMADS27 module"

### Supplemental information for "OsNLP4-OsMADS27 module controls nitrate-dependent salt tolerance in rice" by Alfatih et al.

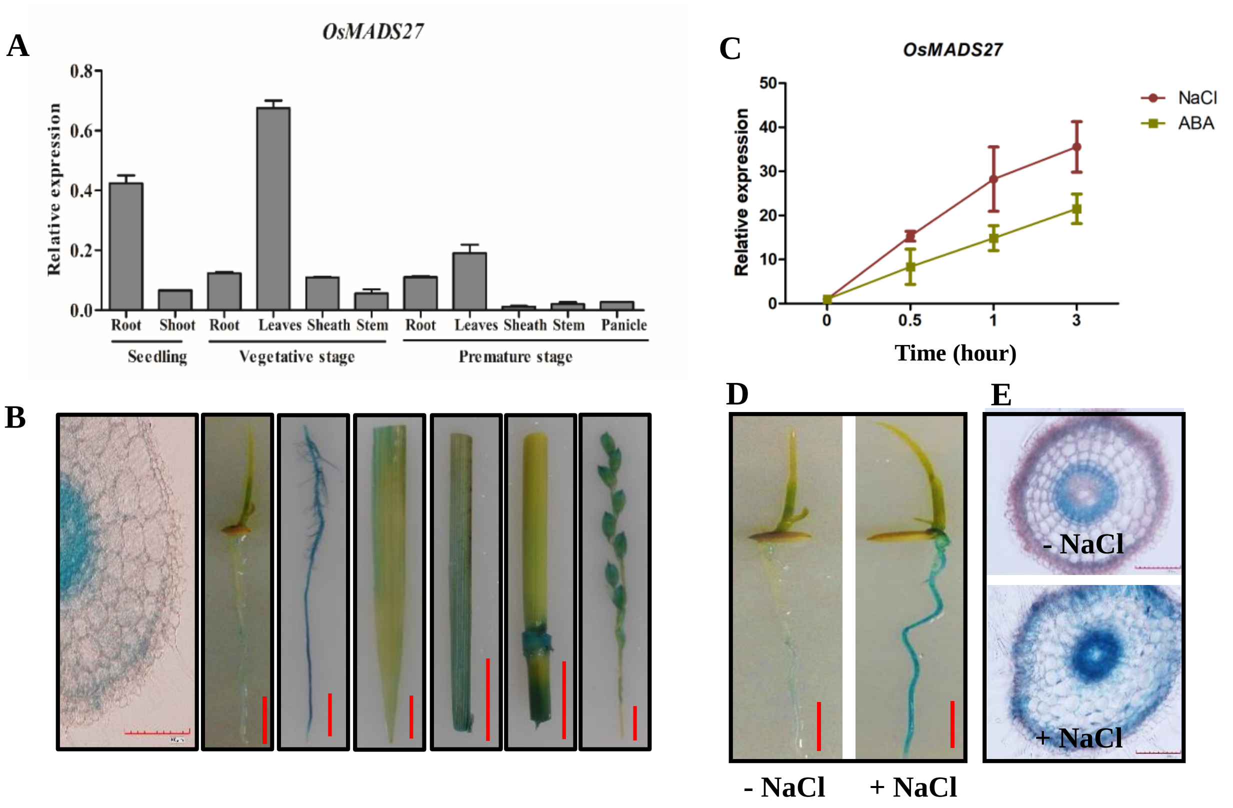


**Figure S1. Expression pattern of *OsMADS27*.**

A. *OsMADS27* transcript levels in different tissues, as determined using qRT-PCR analysis. Values are mean ± SD (n = 3).

B. Expression pattern of *OsMADS27pro:GUS* in transgenic lines. 1-week-old seedling root cross section (Bar = 100 µm), 1-week-old seedling root, 3-month-old plant root, leaf, sheath, stem, and panicle, respectively. Bar = 1 cm.

C. *OsMADS27* is induced by NaCl and ABA. 7-day-old wild type plants grown on normal medium with 140 mM NaCl or 0.01 mM ABA for 0, 0.5, 1, 3 hours, then *OsMADS27* expression levels were detected by qRT-PCR. Values are the mean ± SD (n=3).

D-E. *OsMADS27pro:GUS* responds to NaCl. 7-day-old *OsMADS27pro:GUS* lines grown on MS medium with or without NaCl treatment for 7 days, then seedlings were incubated in GUS buffer for 3 h. The photographs of seedlings (D) and root cross sections (E) were taken. Bar = 1 cm.

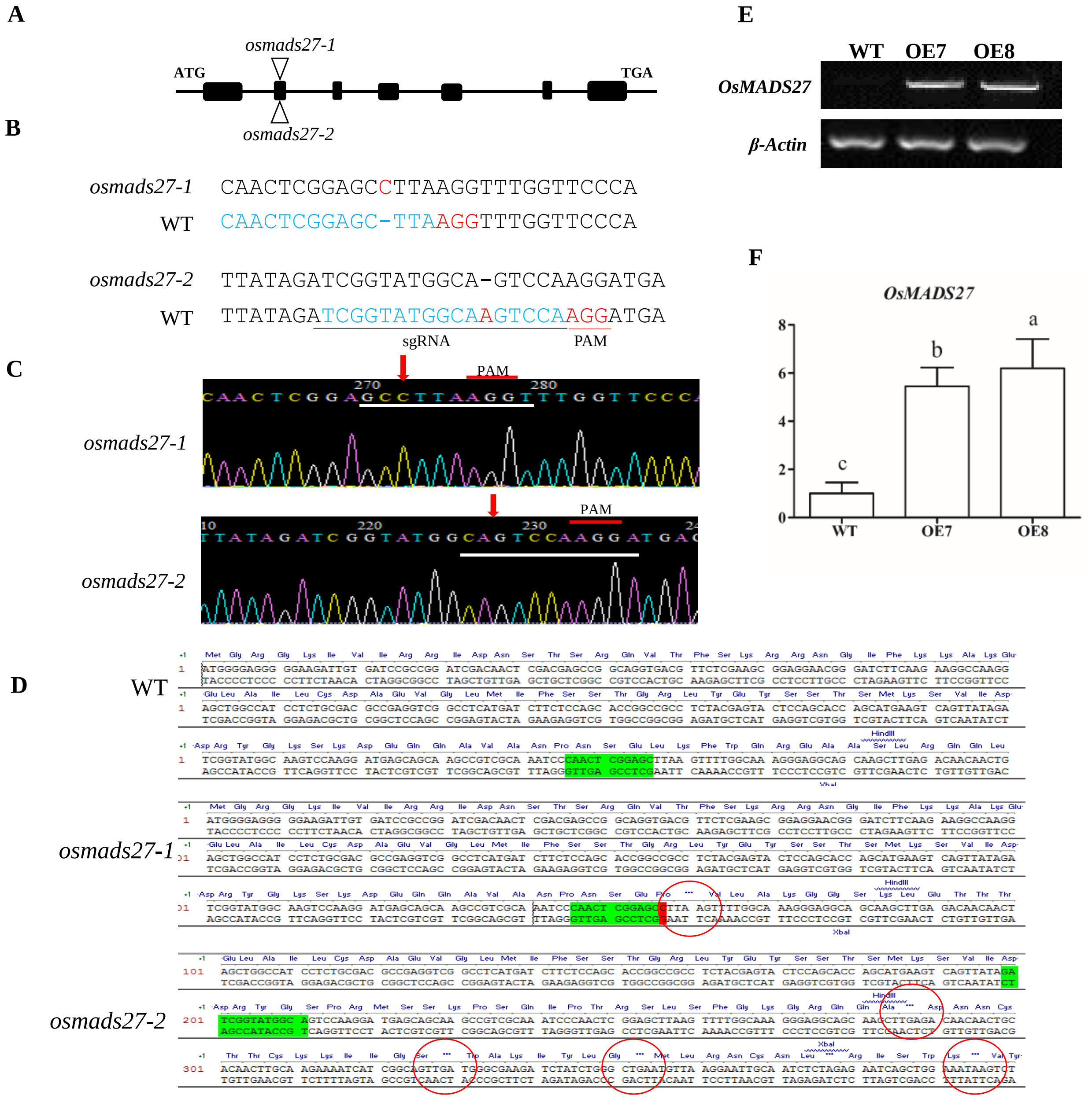


**Figure S2. Verification of the CRISPR/Cas9-edited mutations in *OsMADS27* and overexpression lines of *OsMADS27*.**

A. Schematic diagram of the CRISPR/Cas9-edited sites in two mutant alleles, named KO1 (*osmads27-1*) and KO2 (*osmads27-2*). Triangles indicate the mutated sites in *OsMADS27*.

B. DNA sequences of the edited sites in *OsMADS27*. The dashed line indicates the deletion in the DNA sequences. sgRNA, single guide RNA; PAM, protospacer adjacent motif.

C. Sequence confirmation of the edited sites in KO1 and KO2 mutants. Arrows indicate the edited sites.

D. Reading frame shift in the KO mutants.

E and F. Expression level of *OsMADS27* in the overexpression lines OE7 and OE8 in comparison with wild type by semi-quantitative PCR (E) and quantitative real-time PCR (F). *OsACTIN1* was used as internal control. Values are the mean ± SD (n=3). Different letters denote significant differences ($P < 0.05$) from Duncan’s multiple range tests.

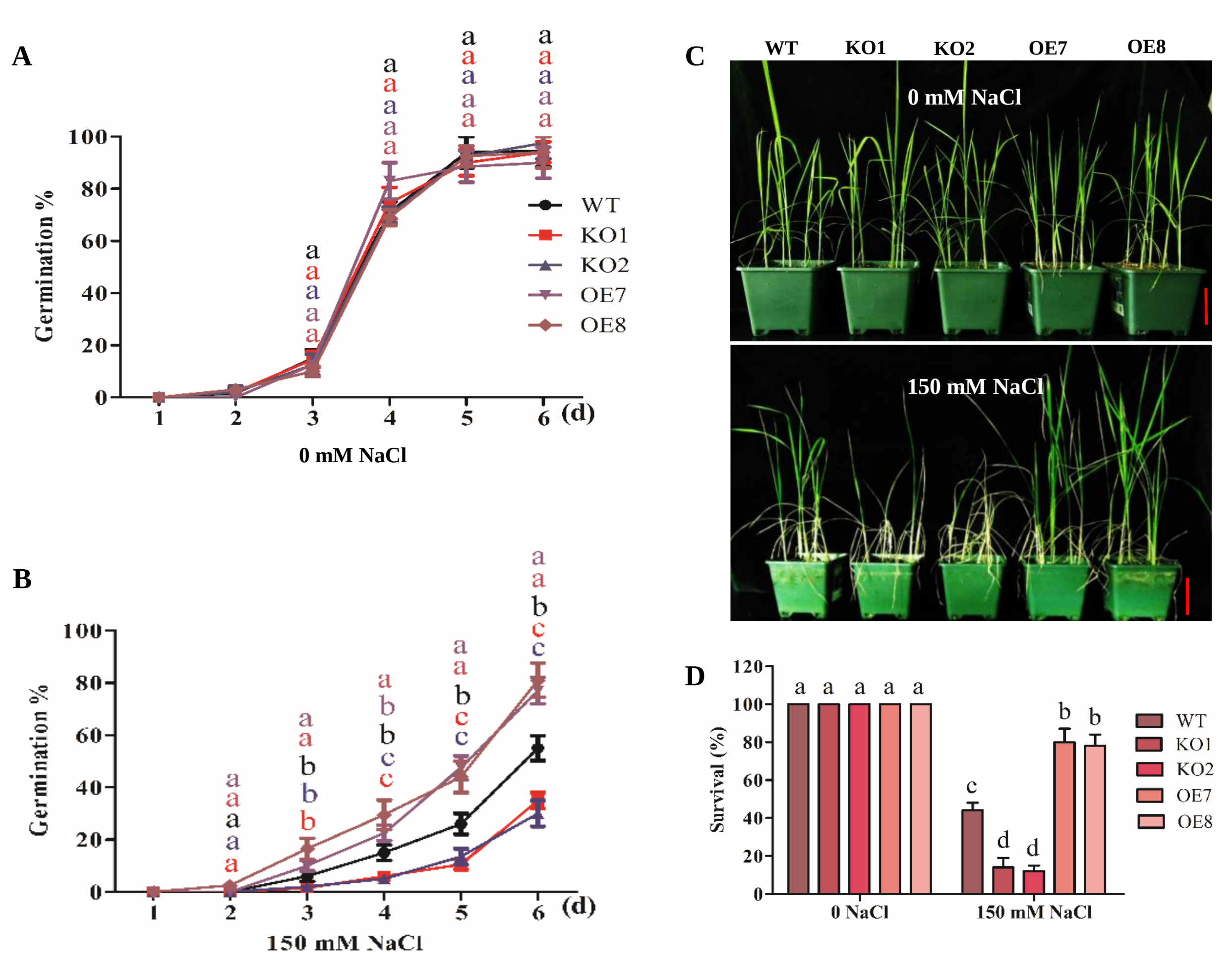


**Figure S3.** ***OsMADS27*** **positively affects salt tolerance in germination and seedling growth.**

A-B. Seed germination of WT, KO1, KO2, OE7, and OE8. Seeds were treated with 0 (A), or 150 mM NaCl (B), and incubated at 37 °C for 7 days. Germination rate was calculated. Values are mean ± SD (n=3 replicates, 60~80 seeds/replicate).

C-D. Salt tolerance assay in soil. 4-week-old soil-grown seedlings of WT (ZH11), KO1, KO2 mutants, OE7 and OE8 lines were irrigated with or without 150 mM NaCl for 2 weeks before photographs were taken (C), and survival rate was calculated (D). Values are mean ± SD (n=6 replicates, 5 seedlings/replicate).

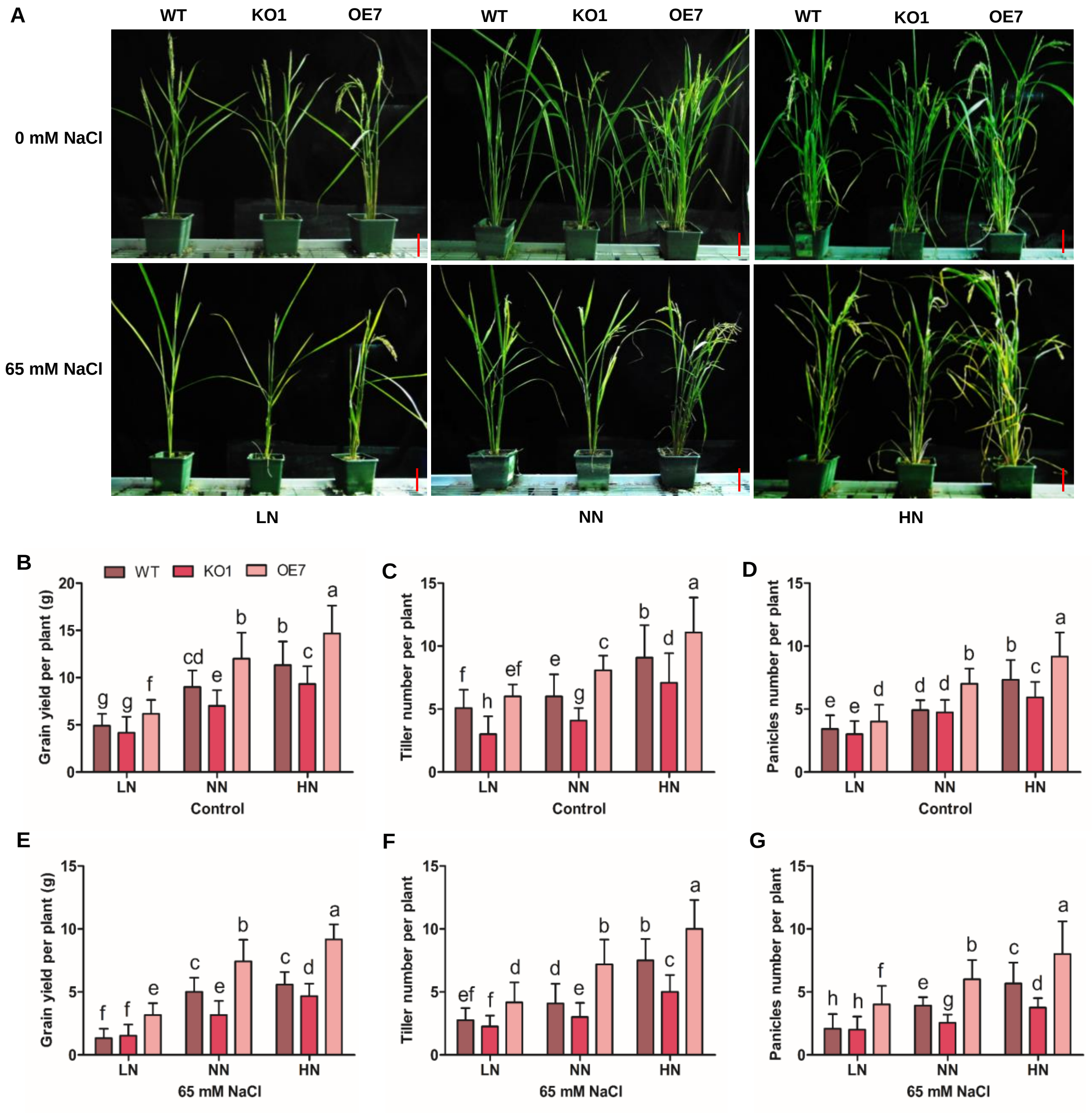


**Figure S4. OsMADS7 positively affects grain yield in a N-dependent manner under normal and salt stress conditions.**

A-G. Salt tolerance assay in vermiculite. 3-week-old vermiculite-grown seedlings of WT, KO1, KO2 mutants, OE7, and OE8 lines were fed with nutrient solutions containing different concentrations of $NO_3$- (1.5 mM LN, 2.5 mM NN, 5 mM HN) and irrigated with or without 65 mM NaCl for 3 times during the rest of growth stages (A). Scale bar = 10 cm. Grain yield per plant (B, E), tillers number per plant (C, F), panicles number per plant (D, G) were calculated, respectively. Values are mean ± SD (n=3 replicates, 16 plants per replicate). Different letters denote significant differences (P < 0.05) from Duncan's multiple range tests.

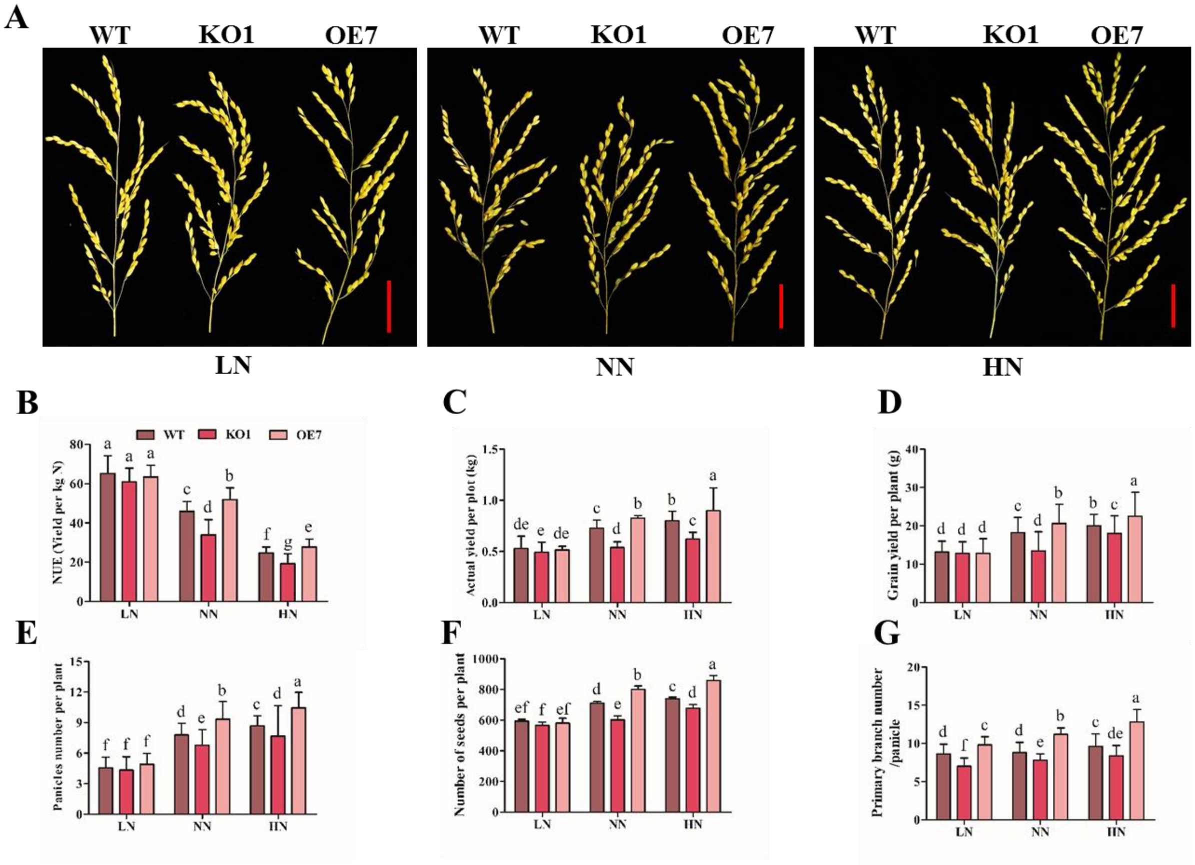


**Figure S5.** ***OsMADS27*** **improves NUE and grain yield in the field of different nitrogen concentrations**.

A. The panicles of wild type ZH11 (WT), KO1, and *OsMADS27*-OE7 plants. Scale bar = 4 cm.

B-G. Nitrogen use efficiency (NUE) (B), actual yield per plot (C), grain yield per plant (D), panicles number per plant (E), number of seeds per plant (F), and primary branch number per panicle (G), were calculated, respectively. Values are mean ± SD (n=3 replicates). Different letters denote significant differences (P < 0.05) from Duncan's multiple range tests.

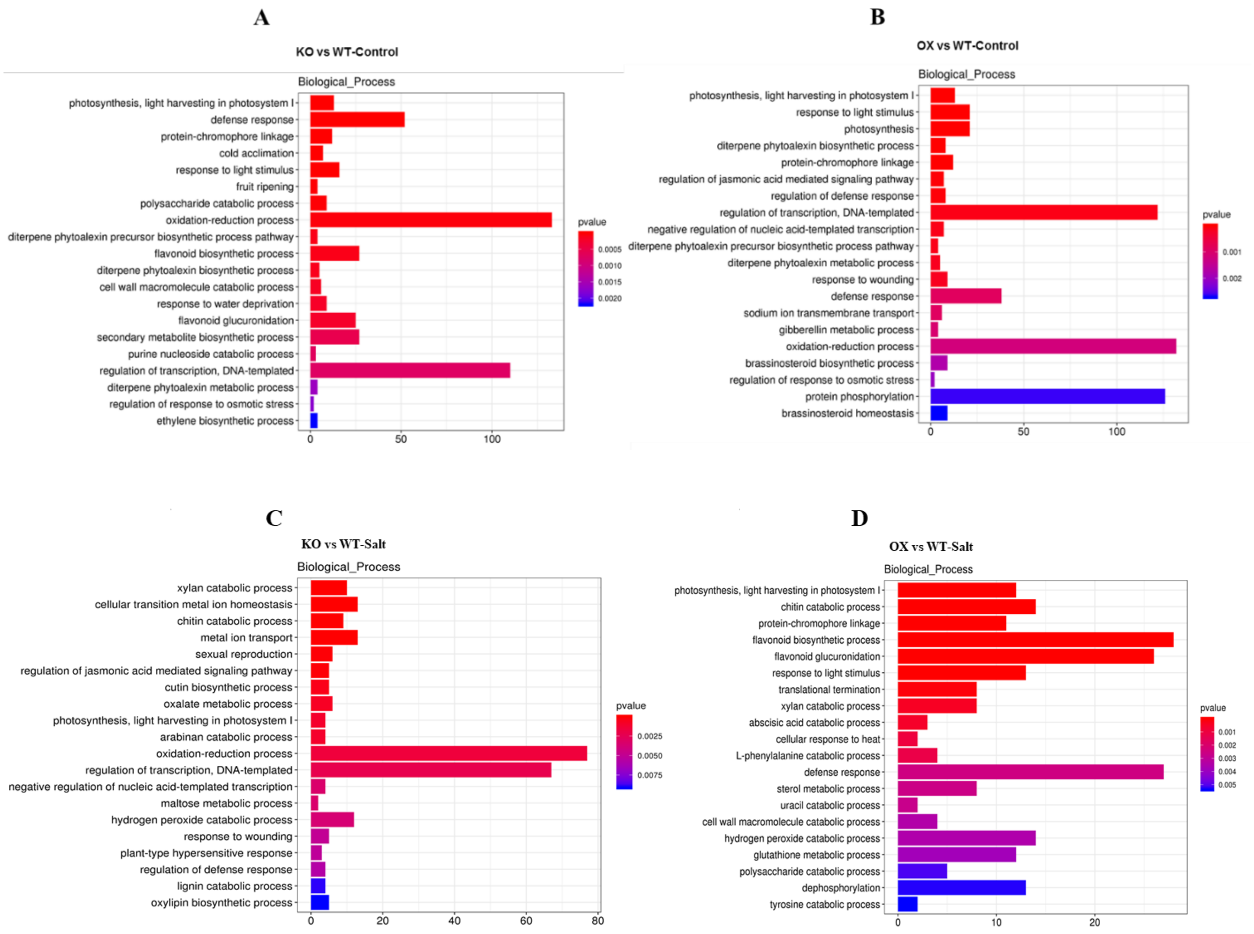


**Figure S6. The gene ontology (GO)-based enrichment analysis of DEGs.**

A-D. number of genes involved in the biological processes in (KO vs WT)-control (A), (OE vs WT)-control (B), (KO vs WT)-salt (C), (OE vs WT)-salt (D).

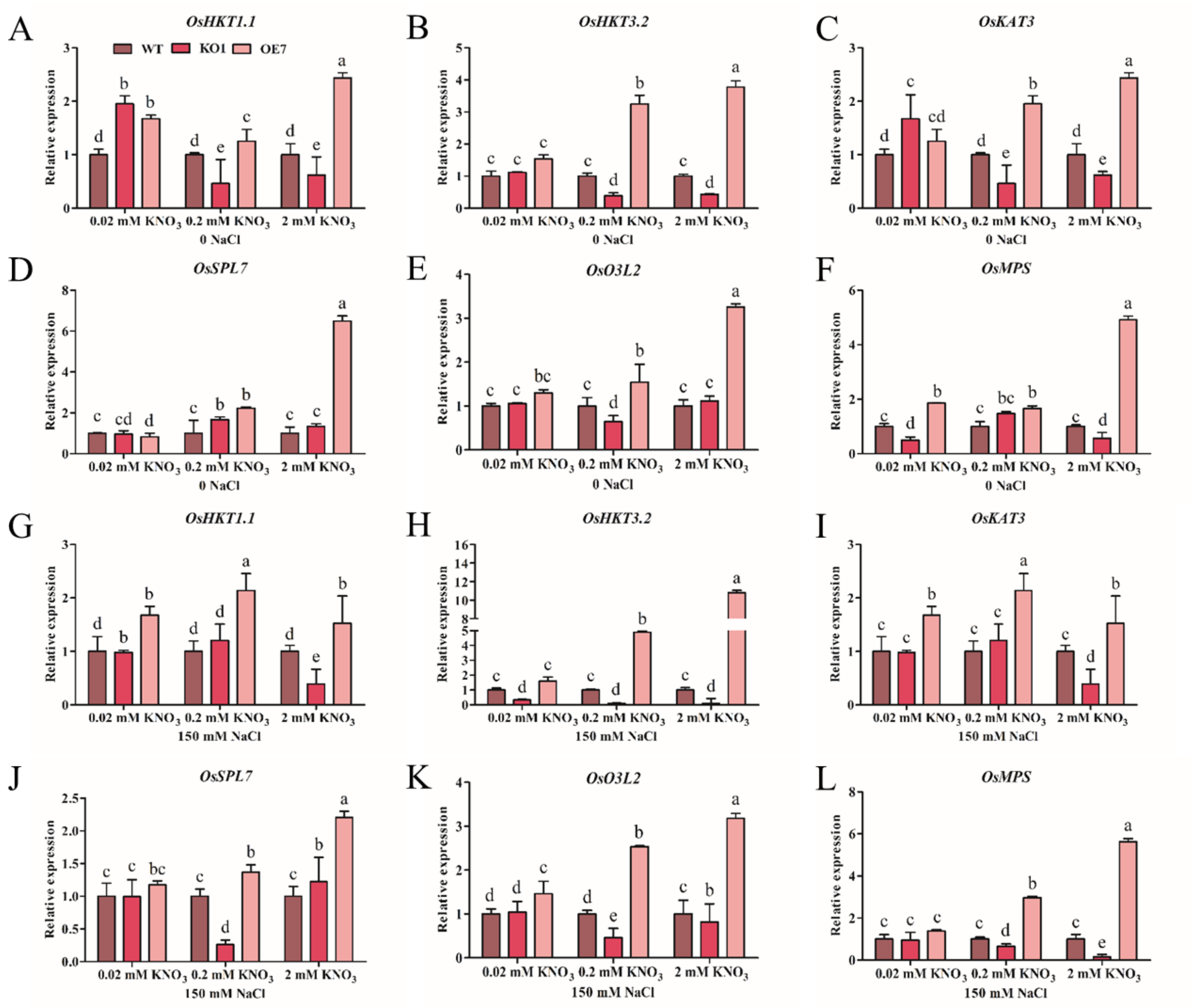


**Figure S7. OsMADS27 broadly regulates the genes involved in salt tolerance.**

7-day-old plants grown hydroponically on medium contain different N concentrations (0.02, 0.2 and 2 mM $KNO_3$) supplemented with either 0 mM NaCl or 150 mM NaCl and then harvested for qRT-PCR analysis. *Actin* was used as an internal control.

A-C. Potassium transporter *OsHKT1.1*, *OsHKT2.3* and *OsKAT3*.

D. Heat shock transcription factor *OsSPL7*.

E. Oxidative stress 3 *O3L2*.

F. R2R3-type MYB transcription factor *OsMPS*.

G-L. Relative expression of the same genes in plants grown on medium with different N concentrations supplemented with 150 mM NaCl. Different letters denote significant differences (P < 0.05) from Duncan's multiple range tests.

**Table S1**. Lists of primers used in this study.

| | |
|---|---|
| ***OsMADS27* related primers** | |
| *OsMADS27*-OE-LP | ATGGGGAGGGGGAAGATTGT |
| *OsMADS27*-OE-RP | TCATGGATTCAACTGTAACCCTA |
| *KO-CRISPR*-LP | TGAAGTGTATAGTGTGTGTCTC |
| *KO-CRISPR*-LP | GGTGTTCTACTGTGTCTATATG |
| *OsMADS27-target1* | AAATCCCAACTCGGAGCTTAAGG |
| *OsMADS27-target2* | AGATCGGTATGGCAAGTCCAAGG |
| *OsMADS27-LP-qPCR* | AGTCAATCGGGATTCACCAA |
| *OsMADS27-RP-qPCR* | AGTTGATTGTTCGGCGTCAC |
| Rice Actin1-LP | TGGCATCTCTCAGCACATTCC |
| Rice Actin1-RP | TGCACAATGGATGGGTCAGA |
| Ubi-*OsMADS27*-GFP-LP | ATGGGGAGGGGGAAGATTGTG |
| Ubi-*OsMADS27*-GFP-RP | TGGATTCAACTGTAACCCTAGC |
| *MADS27pro-OsMADS27*-GFP-LP | GAGCGAGGAGGAAGAACGGA |
| *MADS27pro -OsMADS27*-GFP-RP | TGGATTCAACTGTAACCTGCAT |
| *OsMADS27*-promoter- GUS-LP | GAGCGAGGAGGAAGAACGGA |
| *OsMADS27*-promoter-GUS-RP | CTTCTTCGCCGCTATCTCGA |
| **Salt related genes primers** | |
| *OsHKT1.1-LP* | ATGCATCCACCAAGTTTAGTGCT |
| *OsHKT1.1-RP* | GATTGTTTGACAGAACTTGCCAC |
| *OsHKT2.3-LP* | ATGCCTATTCGGCTGCATATCT |
| *OsHKT2.3-RP* | GAGGATTGTACTTGTGGTTGCT |
| *OsKAT3-LP* | ATCTTGCTTCCACCAGTTCTGG |
| *OsKAT3-RP* | ACTAGGACTATAAGGAACAGCT |
| *OsSPL7-LP* | ATGGAGAGTTCCAACCTGGGCG |
| *OsSPL7-RP* | GAGAAGTTGTTGTGCTTGAAG |
| *OsO3L2-LP* | GGAGATCAAGGTGGTGTTGCTG |
| *OsO3L2-RP* | ACGAGCAGTAGTCACCACCAC |
| *OsMPS-LP* | ATGGAAGGGCAGCAGTTCGCTT |
| *OsMPS-RP* | TCAGGTCTCAGGTAGTTCACCC |
| **ChIP related genes primers** | |
| ChIP- HKT1.1- control-LP | GGATTTCTTACTTATTTATCAGAGT |
| ChIP- HKT1.1- control-RP | GAAACTTTAGGGACTAAAGTAGC |
| ChIP1-HKT1.1-LP | TGGCAAAATGCTGGTTGAACAG |
| ChIP1-HKT1.1-RP | CTTTAACCTGTTTTTTCCACAGC |

| | |
|---|---|
| ChIP2-HKT1.1-LP | GCTGTGGAAAAAACAGGTTAAAG |
| ChIP2-HKT1.1-RP | ACTCTGATAAATAAGTAAGAAATCC |
| ChIP- MADS27- control-LP | TTCGATTTCTATTGAGTAGAGGTT |
| ChIP- MADS27- control-RP | GTTGTACTAGATGATGTCTCATC |
| ChIP1-MADS27-Cis1-LP | GATGAGACATCATCTAGTACAAC |
| ChIP1-MADS27-Cis1-RP | AAGAGAGGAAAGAGGAGACAGC |
| ChIP2-MADS27-Cis2-LP | GCTGTCTCCTCTTTCCTCTCTT |
| ChIP2-MADS27-Cis2-RP | GGATCTAGCTGGGAGCTGGTA |
| ChIP3-MADS27-Cis3-LP | TACCAGCTCCCAGCTAGATCC |
| ChIP3-MADS27-Cis3-RP | CTTCTTCGCCGCTATCTCGATC |
| ChIP- SPL7- control-LP | AGATTAGATTATCTTAGACATC |
| ChIP- SPL7- control-RP | CGATAAAACCCTAATTGGCAAT |
| ChIP- SPL7- Cis1-LP | AAGACGATTGGTCAAACAGTGC |
| ChIP- SPL7- Cis1-RP | CTACTTTACTTGTGACTTATGT |
| ChIP- SPL7- Cis2-LP | GATGACAACAAAGGTTTCACTG |
| ChIP- SPL7- Cis2-RP | GCAATCGGCTGATGATGATGTA |
| ChIP- SPL7- Cis3-LP | ACTTAGTGCTGCATTAGTTAAG |
| ChIP- SPL7- Cis3-RP | CAGTGAAACCTTTGTTGTCATC |
| **Yeast one hybrid related genes primers** | |
| OsMADS27-AD-LP | ATGGGGAGGGGGAAGATTGT |
| OsMADS27-AD-RP | TCATGGATTCAACTGTAACCCTA |
| pHIS2-*HKT1.1*-LP1 | CCTAACCTTTTAAATGGTCTAAATA |
| pHIS2- *HKT1.1*-RP1 | TATTTAGACCATTTAAAAGGTTAGG |
| pHIS2-*SPL7*-Cis1-LP | CTCCCTCCGTCCCAAAATAAGTGCAGA |
| pHIS2-*SPL7*- Cis1-RP | CGCGTCTGCACTTATTTTGGGACGGAGGGAGAGCT |
| pHIS2-*SPL7*- Cis2-LP | CAACCTTCGGATTAATGCTAATTCTAA |
| pHIS2-*SPL7*- Cis2-RP | CGCGTTAGAATTAGCATTAATCCGAAGGTTGAGCT |
| OsNLP4-AD-LP | CGGCTAGCATGGAAGAGGGAGACCCCCAGCCCA |
| OsNLP4-AD-RP | CCCTCGAGTCATGAGAAACCAGTGTGACCAAT |
| pHIS2-*MADS27*-Cis1-LP | CTCATGTCTCATCCTCACTAAGGTTGTTTTTTTTGGGACAAATGGAA |
| pHIS2-*MADS27*-Cis1-RP | CGCGTTCCATTTGTCCCAAAAAAAACAACCTTAGTGAGGATGAGACATGAGAGCT |
| **LUC related genes** | |
| pRI101-MADS27-KpnI-LP | CGGGGTACCATGTACCCATACGATGTTCCAGATTACGCTATGGGGAGGGGGAAGATTGT |
| pRI101-MADS27-KpnI-RP | CCGGAATTCTCATGGATTCAACTGTAACCCTA |
| pGreen-HKT1.1-SalI-LP | GGGCCCCCCCTCGAGGTCGACTGGTCTCTTCGGATTATAGCCA |

| | |
|---|---|
| pGreen-HKT1.1-SalI-RP | CGCTCTAGAACTAGTGGATCCGGTATTGGAGTTGGCAACCCA |
| pGreen-SPL7-LP | GGGCCCCCCCTCGAGGTCGACGATCTACTAAGTCGTGCTTAGAA |
| pGreen-SPL7-RP | CGCTCTAGAACTAGTGGATCCCATGAATACTACTTTACTTGTGAC |
| pRI101-NLP4-LP | TCCCCCGGGATGGAAGAGGGAGACCCCCAGCCCA |
| pRI101-NLP4-RP | GGGGTACCTCATGAGAAACCAGTGTGACCAAT |
| pGreen-MADS27-LP | CCCTCGAGGTGAGGATACAAAAATAGTGAAATA |
| pGreen-MADS27-RP | GCGTCGACTCTTGATCTGAGTATCAACTCTTGA |
| **EMSA related genes** | |
| MBP-OsNLP4-LP | ATCGAGGGAAGGATTTCAGAATTCATGGAAGAGGGAGACCCCCAGCCCA |
| MBP-OsNLP4-RP | CAAGCTTGCCTGCAGGTCGACTCTAGATCATGAGAAACCAGTGTGACCAAT |
| OsMADS27-cis1+LP | 5'biotinTCATGTCTCATCCTCACTAAGGTTGTTTTTTTTGGGACAAATGGA3' |
| OsMADS27-cis1+RP | TCCATTTGTCCCAAAAAAAACAACCTTAGTGAGGATGAGACATGA |
| OsMADS27-cis1-LP | TCATGTCTCATCCTCACTAAGGTTGTTTTTTTTGGGACAAATGGA |
